# Elucidating the unknown transcriptional responses and PHR1 mediated biotic and abiotic stress tolerance during phosphorus-limitation

**DOI:** 10.1101/2022.08.16.504161

**Authors:** Wolf-Rűdiger Scheible, Pooja Pandey-Pant, Bikram D. Pant, Nick Krom, Randy D. Allen, Kirankumar S. Mysore

## Abstract

Phosphorus (P) limitation in the majority of world soils is a major constraint for plant growth and crop productivity. RNA sequencing was used to discover novel P-responsive gene transcripts (PRGT) in leaves and roots of Arabidopsis. Hisat StringTie and Cufflinks TopHat transcript assembler were used to analyze reads and identify 1,074 PRGTs with a >5-fold altered abundance during P-limitation. Interestingly, 60% of these transcripts were not previously reported. Among the novel PRGT, 106 were from unannotated genes, and some were among the most P-responsive, including *At2g36727* which encodes a novel microRNA. Annotated novel PRGTs encode for transcription factors, microRNAs, small signaling peptides, long non-coding RNAs, defense-related proteins, and transporters, along with proteins involved in many biological processes. We identified several genes that undergo alternative splicing during P-limitation, including a novel *miR399* resistant splice variant of *PHOSPHATE2* (*PHO2*.*2*). Several novel P-responsive genes were regulated by *PHOSPHATE STARVATION RESPONSE1* (*PHR1*), *PHR1-LIKE 1* (*PHL1*) and *PHO2*. We discovered that P-limited plants show increased resistance to pathogens and drought stress mediated by PHR1-PHL1. Identification of novel P-responsive transcripts and the discovery of the influence of P-limitation on biotic and abiotic stress adds a significant component to our understanding of plant P-signaling.

**Highlight:** Phosphorus limitation elicits the expression of several novel genes including many previously unannotated genes, noncoding RNAs, small peptides and alternatively spliced RNAs, and leads to enhanced disease and drought tolerance.

## INTRODUCTION

Phosphorus (P) is an essential element required for plant growth. P is an indispensable component of cellular intermediates in central and energy metabolism, signaling molecules, phospholipids and nucleic acids (Stigter and Plaxton, 2015). P is often the most limiting element for plant growth and development (Holford, 1997; Darcy et al., 2018), and application of P-fertilizers is required to enhance soil P availability and optimize crop yields. However, P-fertilizer use poses a sustainability challenge. Intensive application of P enables high-yield agriculture but also contributes to eutrophication of aquatic systems, while inadequate access to affordable P fertilizer limits crop yields in many developing countries. Moreover, mined rock P, the primary source of P-fertilizer, is a finite resource subject to large price fluctuations and geopolitical risks (Cordell and White, 2014). P sustainability is vital for food security and improvements in plant P efficiency are required through better acquisition or use with the ultimate goal of crop plant species that tolerate lower P availability without reduction of biomass yield or quality. Informed approaches to develop such plants build upon a system-wide understanding of how plants react, respond, and adjust to P limitation.

P limitation responses in plants are drastic and occur at different levels (Fredeen et al., 1989; Robinson, 1994; Lynch, 1995; Darcy et al., 2018; Hou et al., 2020). Among these, the transcriptional response may be the most amenable. Using hybridization-based technologies for transcript profiling, such as Affymetrix ATH1 and tiling gene chips encompassing probe sets for over 22,000 genes, steady state levels of many gene transcripts were reported to change significantly during P-limitation in *Arabidopsis thaliana* (Misson et al., 2005; Morcuende et al., 2007; Bustos et al., 2010; Woo et al., 2012). Similar transcriptome studies were also done in rice (Li et al., 2010; Secco et al., 2013) and soybean (Wang et al., 2016). The three ATH1 gene chip studies by Misson et al., Morcuende et al. and Bustos et al. are highly cited and provide benchmarks for *Arabidopsis*. These studies revealed major insights in a plant species’ response to P limitation/deprivation, as the functional classification of P-status responsive gene transcripts (PRGTs) indicated adaptations of (i) uptake and transport of Pi and other inorganic ions (sulfate, iron), (ii) Pi salvage systems, (iii) lipid biosynthetic pathways, (iv) primary and secondary metabolism, (v) phytohormone synthesis/response pathways, (vi) cell wall metabolism and (vii) transcriptional and posttranslational signaling.

Despite such major insight, the readout from ATH1 gene chips remained incomplete due to partial and biased gene representation, and limited sensitivity. For comparison, The Arabidopsis Information Resource (TAIR10) genome annotation contains 33,602 genes i.e. roughly 50% more than what is represented in ATH1 Gene chips. Dedicated and highly sensitive approaches such as quantitative reverse-transcription PCR (RT-qPCR) revealed additional information about the transcriptional P-starvation response in *Arabidopsis*. For example, Morcuende et al. (2007) profiled ∼2200 transcription factor genes and other gene families (e.g., phosphate transporters, purple acid phosphatases, PHO1-like and SPX-domain proteins, phospholipases and GDPDs) for their response to P-limitation, revealing P-responsive genes not found or represented on ATH1 arrays. Similarly, Pant et al. (2009) analyzed the expression of ∼200 microRNA primary transcripts, none of which were found on the ATH1 array. This study confirmed the strong induction of several miR399 species (Bari et al., 2006; Pant et al., 2008), and also revealed strong induction of *miR778, miR827* or *miR2111* during P-limitation.

The advent of high-throughput RNA sequencing (RNA-seq) technology provided the opportunity to reanalyze the P-starvation response of *Arabidopsis* and other plants at truly genome-wide depth, without bias and with sensitivity limited only by the number of RNA-seq reads produced. As yet, no dedicated attempt has been made to comprehensively describe the transcriptional response to P-starvation response in the model plant *Arabidopsis* with this technology. A major obstacle for doing so may be related to bioinformatics challenges, including the availability of software for read mapping and proper transcript assembly (Pertea et al., 2015) to ensure accurate reconstruction of a transcriptome from the RNA-seq reads. Genome-wide DNA methylation patterns in response to phosphate starvation were determined and correlated with the RNA-seq-based expression levels of P-starvation responsive genes (Yong-Villalobos et al., 2015). However, this study lacked information about the unknown transcriptional P-starvation response. RNA-seq was also performed to compare the response of P-limitation in roots of *Arabidopsis* wild-type and *PHOSPHATE RESPONSE1* (*PHR1*)-*LIKE 2* (*phl2*) mutant (Sun et al., 2016). This analysis provided a broad comparison of 581 P-starvation-upregulated (≥2-fold) gene transcripts in the two genotypes at the gene ontology level. The result showed a >30% reduction in expression of a majority (88%) of these phosphate starvation-inducible (PSI) genes in the *phl2* mutant. A third study (Yuan et al., 2016) mined RNA-seq data from P-starved *Arabidopsis thaliana* for long non-coding RNAs with altered abundance but information about other gene classes or novel protein-coding PRGTs was not reported.

To help fill this void, we analyzed RNA-seq data from roots and shoots of P-sufficient and P-limited *Arabidopsis* plants, grown in sand culture, to identify previously unknown P-starvation responsive gene transcripts in this model plant species. We also compared the obtained expression data with the ATH1 results reported by Bustos et al. (2010), because this study, provided data for both shoots and roots and for all genes represented on the gene chip. We analyzed novel *PSI* genes for their reversible induction to P-availability by RT-qPCR, their regulation by *PHR1/PHL1* and *PHOSPHATE2* (*PHO2*), and screened for P-responsive alternative splicing forms. We analyzed the novel PRGTs for their involvement in different biological processes and cross-talk with other biotic and abiotic stresses and we analyzed P-limited *Arabidopsis* plants for resistance to a bacterial pathogen and drought stress.

## MATERIALS & METHODS

### Plant materials and nutrient media

*Arabidopsis thaliana* Columbia-0 wild-type (WT) seeds were surface sterilized and grown in half strength Murashige and Skoog (MS) nutrient agar plates for 10 days under short-day conditions with light/dark regime of 8/16 hours in a Percival growth chamber (www.percival-scientific.com). Seedlings were transferred to acid-washed sand and perlite mixture and grown in a growth chamber with a light/dark regime of 12/12 hours at 23/22 ºC, respectively, with a light intensity of ∼125 µmol m^-2^ s^-1^ and a relative humidity of 50%. The plants were divided into two groups and were watered with the nutrient solution with full nutrition (P-sufficient, +P) and without phosphate (-P) for the first three days and then the nutrient solution was diluted 5X and supplied to the plants every other day. Plants were grown under -P condition for nine days until the typical P-limitation phenotype (dark-green leaves and smaller plants) appeared. The location of the plants within a growth chamber were randomized every two days to minimize position effects. Pi-limitation in plants was confirmed before harvesting by measuring inorganic phosphate in the leaves. The roots and shoots were washed with deionized water, harvested, snap frozen in liquid nitrogen and stored at -80 ºC until further analysis. The nutrient media contained 4 mM KNO_3_, 2 mM NH_4_NO_3_, 3 mM KH_2_PO_4_/K_2_HPO_4_, *pH* 5.7, 4 mM CaCl_2_, 1 mM MgSO_4_, 2mM K_2_SO_4_, 1 mM MES, *pH* 5.8 (KOH) and microelements/nutrients: ZnSO_4_ (1 μM), Na_2_FeEDTA (40 μM), H_3_BO_3_ (60μM), MnSO_4_ (14μM), CuSO_4_ (0.6 μM), NiCl_2_ (0.4μM), HMoO_4_ (0.3 μM), CoCl_2_ (20 nM). The –P media contained no KH_2_PO_4_/K_2_HPO_4_ but 2.5 mM KCl was added instead. For the phosphate re-addition experiment, the plants grown under +P and –P conditions were re-supplied with 600 µM KH_2_PO_4_/K_2_HPO_4_-containing nutrient media and harvested after 3 and 6 hours. *Arabidopsis* WT, *phr1, phr1phl1* and *pho2* mutants (Rubio et al., 2001; Bari et al., 2006; Bustos et al., 2010) were grown in axenic liquid culture as described previously (Pant et al., 2015).

### Phosphate (Pi) measurement

Measurement of the inorganic phosphate was done using the colorimetric micro method (Itaya and Ui, 1966) as described in Pant et al. (2015). In summary, frozen plant material was ground in liquid nitrogen, weighed and soluble inorganic phosphate was extracted in Milli-Q water (Millipore) by repeatedly (3X) freezing and heating the samples at 95 °C for 5 minutes. 10μL of the sample was mixed with 100μL HCl and 100μL malachite green color reagent in a 96 well plate. The plate was incubated at room temperature for 15 min. and the absorbance was measured at 630 nm. Phosphate concentration in the samples was determined based on a calibration curve.

### RNA isolation, cDNA synthesis and quantitative reverse transcription PCR

RNA was isolated using TRIzol reagent (Invitrogen) according to the manufacturer’s standard protocol. For RT-qPCR and RNA-seq experiments, residual DNA was removed from RNA samples by treating with TURBO DNA-free™ DNase (Ambion) according to manufacturer’s instructions. The complete removal of genomic DNA (gDNA) from RNA was confirmed by RT-qPCR using the intron or exon primers, where gDNA was taken as a positive control and water sample as a negative control. NanoDrop spectrophotometer (ND-8000) was used to check the concentration and purity of RNA, and Agilent Bioanalyzer 2100 was used to check the RNA integrity. RNA samples with RNA Integrity Number (RIN) values >8 were taken for RNA-seq. Selection of primers, RT-qPCR conditions and procedures for cDNA synthesis were as described previously (Pant et al., 2008; Pant et al., 2015). DNA oligos used in the study are given in supplemental table S8.

### RNA-seq library preparation and deep sequencing

Three independently prepared batches of plants grown under P-sufficient and P-limited condition were used for RNA-seq experiments. Materials from five plants were pooled for each of the root and shoot samples. RNA-seq libraries were prepared at the genomics core facility of the Noble Research Institute using the Illumina® TruSeq® RNA Sample Preparation Kit v2, according to manufacturer’s instructions. This includes mRNA isolation and purification from total RNA using oligo dT magnetic beads. The Illumina TruSeq kit v2 set “A” indexed adapter sequences were ligated to each sample for multiplexed sequencing. Quality control of the RNA and cDNA libraries was performed using Agilent 2100 Bioanalyzer (Agilent Technologies). RNA-seq libraries were sequenced on a flow cell of Illumina HiSeq2000 sequencing instrument.

### RNA-seq read mapping and data processing

Paired-end reads generated from the RNA-seq were used for each sample. All RNA-seq reads were demultiplexed using a known list of barcodes (Illumina), allowing zero mismatches. After demultiplexing, adapter sequences were removed from the sequenced libraries using the FastQ/A clipper found in the FastX Toolkit (http://hannonlab.cshl.edu/fastx_toolkit/). Reads were quality trimmed by removing low-quality bases until two consecutive bases with quality scores of 30 or above were found. Only the quality trimmed reads longer than 50 nucleotides were used for alignment with the *Arabidopsis* reference genome version 10 (TAIR10) using the short read aligner Bowtie 2 V2.2.3.0 and splice junction mapper TopHat2 v2.0.9 (Kim et al., 2013) with default parameters. The mapped reads were used for transcript assembly and quantification using the default parameters in Cufflinks V2.1.1 (Trapnell et al., 2010; Trapnell et al., 2014). Gene and transcript expression levels were normalized by Cufflinks as fragments per kilobase of transcript per million mapped reads (FPKM). A unified set of transcripts was constructed by comparing all the transcripts assembled from each sample using Cuffcompare (Trapnell et al., 2014). Fold change values (-P vs. FN) were calculated by adding 0.1 to each FPKM value, to avoid divisions by zero. To identify PRGTs, the following filtering procedure was performed: shoot PRGTs had to display a ≥ 5-fold average change for the two biological replicates with each replicate being at least 3-fold changed (average [R1+R2]/2 ≥5; R1 ≥3 and R2 ≥3) or (average [R1+R2]/2 ≤0.2; R1<0.333 and R2 ≤0.333). Root PRGTs were selected if the average change for two replicates was at least 5-fold with each of the two replicates showing an at least 3-fold change, and the third replicate showing at least a 2-fold change (average [R1+R2]/2 ≥5; R1 ≥3 and R2 ≥3 and R3 ≥2) or (average [R1+R2]/2 ≤0.2; R1<0.333 and R2 ≤0.333 and R3 ≤0.5). Existence and P-limitation response of the known and unannotated novel genes was confirmed by visualizing the read alignment (Binary Alignment/Map, BAM files) in integrative genomics viewer (IGV version 2.3.72; http://software.broadinstitute.org; (Thorvaldsdottir et al., 2013) and performing RT-qPCR expression analysis for some selected genes/loci.

To obtain more inclusive list of differentially regulated genes in response to P-limitation, the RNA-seq reads were also analyzed using an independent method. Illumina adapters and low-quality bases were trimmed using Trimmomatic (Bolger et al., 2014) using the default parameters: LEADING: 3 TRAILING: 3 SLIDINGWINDOW: 4:15 MINLEN: 36. Quality trimmed reads were mapped to the *Arabidopsis* reference genome to produce Sequence Alignment Map (SAM) files using the HISAT alignment software with the alignment option: -dta-cufflinks (Kim et al., 2015). SAM files were converted into compressed and sorted BAM files using Samtools - view, -bSh, and -sort-sort commands, respectively (Li et al., 2009). Reads mapping to annotated and unannotated regions of the *Arabidopsis* genome were extracted using StringTie (Pertea et al., 2015). Genes with less than one read count per million across at least three samples were discarded from the analysis. For the differential expression analyses, significant transcripts were converted to the corresponding gene for performance evaluation, such that if a single transcript was called as differentially expressed, the corresponding gene was also called differentially expressed. We note that because of this unavoidable difference between gene-level and transcript-level comparisons, quantitative comparisons of recall and/or precision between a gene-level and a transcript-level workflow should be avoided. Rather, we recommend evaluating the relative performance of a given workflow as compared with other workflows with matched gene-level or transcript-level estimation.

### Water Loss Assay

For drought tolerance analysis, water loss assay was performed as described previously (Pant et al., 2022). Briefly, *Arabidopsis* WT and *phr1* mutants were germinated on ½ MS media with vitamins (M519, PhytoTechnology Laboratories, KS, USA) buffered to pH 5.7 with 150 mg/L MES, supplemented with 0.3% (w/v) Phytagel (sigmaaldrich.com), and 1% (w/v) sucrose. Five-day-old plants were transplanted to +P and -P media plates containing ½ MS medium without NPK (PhytoTechnology Laboratories, KS, USA) buffered to pH 5.7 with 150 mg/l MES, supplemented with 1% (w/v) sucrose, 0.3% (w/v) Phytagel (Sigma), 1.5 mM NH_4_NO_3_, 3 mM KNO_3_, 3 mM KH_2_PO_4_/K_2_HPO_4_ (for +P) and 2.5 mM KCl (for -P). The plants were grown on a growth chamber under short-day condition (∼130 μmol m^−2^ s^−1^, 8-h light/16-h dark photoperiod), at 23 °C and 40% relative humidity. Rosette from the 3-week-old plants were detached and allowed to lose water at 23 °C under light (100 µmol m^−2^s^−1^). Weight of the rosette was measured every 1 h for five times before calculating the relative water loss.

### Pathogen Assay

*Arabidopsis thaliana* ecotype Columbia (Col-0) plants were grown in GS90 soil in the Percival AR36-L growth chambers (www.percival-scientific.com) under 8-h light (∼ 150 µmol m^-2^ s^-1^, 24 °C /16-h dark (20 °C) cycle at ∼ 70% relative humidity. *Pseudomonas syringae* pv. *tomato* DC3000 were grown in King’s B medium at 28 °C supplemented with rifampicin (10 mg/mL). Twenty-five days old plants were spray inoculated with *P. syringae* pv. tomato DC3000 at concentration of 1 × 10^8^ CFU/ml in water containing 0.01% Silwet L-77 and the plant phenotype was recorded after 5 days. Pathogen assay for *Erwinia carotovora* ssp. *carotovora* and *P. syringae* pv. *maculicola* was performed as described previously (Ishiga et al., 2011; Pant et al., 2020). Briefly, 3-week-old *Arabidopsis* plants grown on ½ MS media plates (+P and -P condition) were flood inoculated at concentration of 1.6 × 10^5^ CFU/ml in water containing 0.01% Silwet L-77. Quantification of bacterial growth was done at 3 dpi for *E. carotovora* and 5 dpi for *P. syringae* pv. *maculicola* by counting colony forming units (CFUs).

## RESULTS

### RNA-seq study identified several known and novel P-limitation responsive gene transcripts (PRGTs)

Plants grown in P-limitation (–P) were distinctly smaller than plants grown in P-sufficient (+P) conditions and showed anthocyanin accumulation (supplemental Figure S1 A, B). Inorganic phosphate content in the shoots of –P grown seedlings was about 10-fold lower than in +P grown seedlings (supplemental Figure S1 A, B) and the levels of three P-limitation marker gene transcripts (*NON-SPECIFIC PHOSPHOLIPASE C5, NPC5* [*AT3G03540*], *Transcription elongation factor SPT5* [*AT2G34210*] and *INDUCED BY PHOSPHATE STARVATION1, IPS1* [*AT3G09922*]) were strongly increased (supplemental Figure S1C). RNA-Seq library production and analysis resulted in 10 RNA-seq files (supplemental data files 1-10). Analysis of the RNA-seq data was performed using HiSat StringTie and the TopHat Cufflinks pipelines using the stringent approach. The output file from the HiSat StringTie pipeline (supplemental Table S1) contains 26,922 gene entries of which 25,942 have an assigned reference gene. Of the 980 gene entries without annotated reference genes 246, 184, 176, 150 and 200 were on chromosomes 1, 2, 3, 4 and 5, respectively, and another 24 were annotated to the chloroplast and mitochondrial genomes. We identified 788 and 536 PRGTs with ≥ 5-fold change in abundance in shoot and roots, respectively (Figure 1A), while 2,057 and 755 showed ≥ 3-fold change, and 5,029 and 1,293 showed ≥ 2-fold change. Among the highly (≥ 5-fold) responsive PRGTs those that are induced are much more abundant than those that are repressed (610 vs 178 in shoots; 477 vs. 59 in roots). In the shoot, 50 PRGTs were determined to be induced by >100-fold (Figure 1B). Among those 50 PRGTs, 28 are represented on ATH1 gene chips and are known PRGTs. The remaining 22 include some additional known PRGTs identified by other approaches (e.g. *IPS1, miRNA399d*), along with several annotated as “unknown protein” or “other RNA” and six that are not annotated in TAIR10 (Figure 1B).

**Figure 1.**
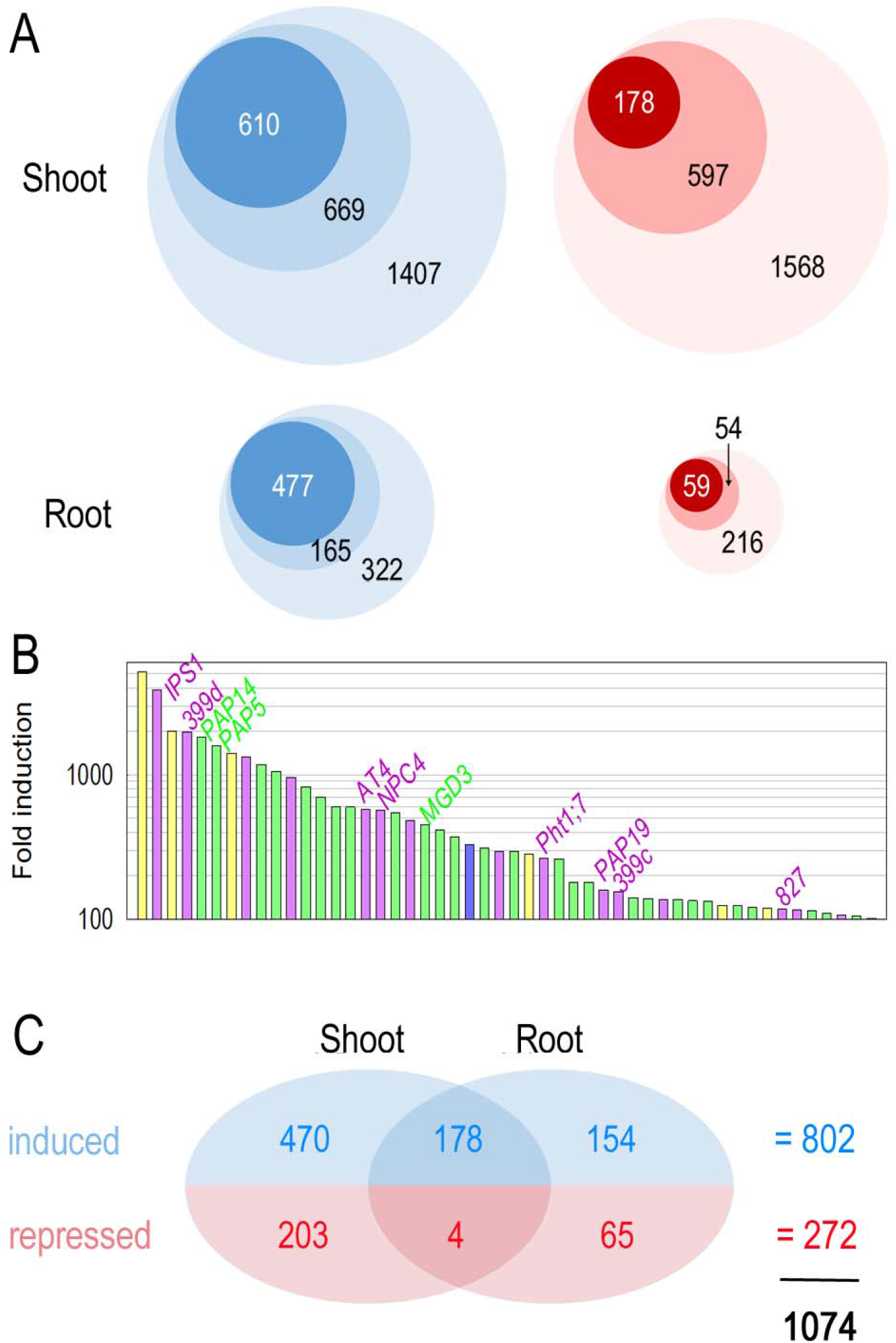
P-status responsive gene transcripts (PRGTs) identified by RNA-seq. (A) Circles indicate the numbers of induced (blue) or repressed (red) gene transcripts in shoot (top) or root (bottom) identified using HiSat StringTie. Circle size is proportional to gene transcript number. Light colors represent ≥2-fold, medium ≥3-fold and darkest colors indicate ≥5-fold change. Refer to Materials and Methods for the definition of a 2-, 3- or 5-fold change. (B) Response of the fifty most strongly induced gene transcripts in shoots. Green and blue bars represent gene transcripts that were also identified in *Arabidopsis* ATH1 gene chip studies. Pink and yellow bars represent PRGTs not present in ATH1 gene chips, with the latter ones also not being annotated in TAIR10 (see legend to Figure 2). (C) Venn diagram showing the numbers of gene transcripts ≥5-fold induced or repressed in shoots, roots or both identified using HiSat StringTie and TopHat Cufflinks analysis. See Table S1 for more details on these altogether 1,074 gene transcripts.

The filtered ≥ 5-fold changed PRGTs were manually inspected for their response in the Integrative Genomics Viewer resulting in 909 and 861 PRGTs in the HISAT-StringTie and TopHat Cufflinks output files, respectively, with an overlap of 696 and a non-redundant total of 1,074 PRGTs (supplemental Table S1, supplemental Figure S2). Reliance on only one of the two RNA-seq analysis pipelines would have precluded the identification of some known PRGTs already identified in ATH1 studies. For example, *SPX DOMAIN GENE 3* (*AT2G45130/ /SPX3*), *AT1G63005/miR399b, S-PHASE KINASE-ASSOCIATED PROTEIN 1*-*LIKE 11* (*AT4G34210*/*ASK11*), *GLYCEROPHOSPHODIESTER PHOSPHODIESTERASE 6* (*AT5G08030/GDPD6*) were identified with HiSat StringTie, whereas *AT2G17280, GALACTINOL SYNTHASE 1* (*AT2G47180/GOLS1*), *MYO-INOSITOL-1-PHOSTPATE SYNTHASE 2* (*AT2G22240/IPS2*) or *AT1G62480* were extracted with the TopHat Cufflinks pipeline only (supplemental Figure S2A). Both analysis pipelines detected 405, 116 and 138 GTs with strong P-response in shoots, in roots or in both shoots and roots, respectively.

Of the 1,074 non-redundant PRGTs, 802 (74.7%) were induced and 272 (25.3%) were repressed during P-limitation (Figure 1C and S1B, supplemental Table S1, categories 1-5). Of the 1,074 PRGTs, 229 >5-fold changed PRGTs detected in this study were reported as >5-fold changed in the ATH1 study of Bustos et al. (2010) (Figure 2A, green sector). In addition, 259 of the 1074 >5-fold changed PRGTs (Figure 2A, pink sector) detected by RNA-seq correspond to TAIR10 annotated genes for which no probe sets are present on ATH1 arrays, and yet another 106 PRGTs (yellow sector) were found for which no gene models exist in TAIR10. Altogether, this RNA-Seq study reports 648 (= 283+259+106; red, pink and yellow sectors respectively) (Figure 2A) strongly differentially expressed PRGTs for which no evidence was previously obtained in Bustos et al. (2010) and the P-responsive status of many was previously unknown. Figure 2B shows Sashimi plots for three PRGTs currently not annotated in TAIR10 (yellow sector), tentatively named *AT5G27993, AT1G53605* and *AT1G53615*.

**Figure 2.**
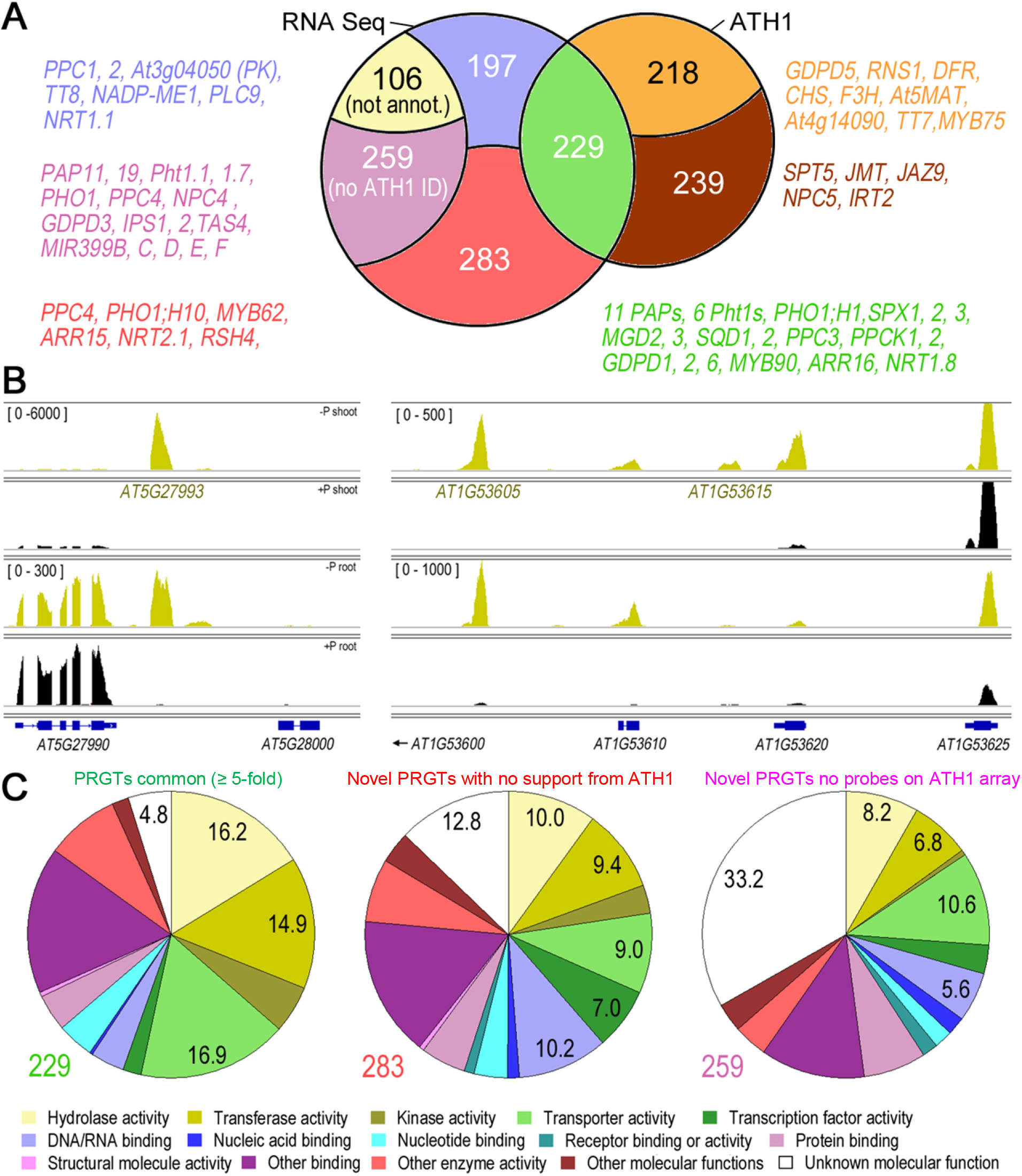
Overview of strong PRGTs identified with RNA-seq. (A) Venn diagram showing the number of PRGTs (5-fold change used as cut-off) identified by RNA-seq and overlap with *Arabidopsis* ATH1 gene chip results from Bustos et al. (2010). The colored sectors are defined as follows: green - GTs ≥5-fold changed in both studies; blue - ≥5-fold changed in RNA-seq and 2-5-fold changed in ATH1 study; red - ≥5-fold changed in RNA-seq but not changed in ATH1 study; pink – GTs ≥5-fold changed in RNA-seq and without probe set on ATH1 array; yellow -GTs of unannotated genes; orange - ≥5-fold changed in ATH1 study and 2-5-fold changed in RNA-seq study; brown - ≥5-fold changed in ATH1 study without support from RNA-seq. Examples of gene transcripts for each sector are given in font with the respective color. (B) Sashimi plots of examples of PRGTs originating from unannotated genes (yellow sector in (A)). Suggested gene codes (*AT5G27993*; *AT1G53605*; *AT1G53615*) are indicated. (C) Gene ontology classification of the PRGTs from the green, red and pink sectors.

RT-qPCR was used to verify the P-limitation response of thirty randomly selected gene transcripts from the green (4), pink (19), yellow (5) blue (1) and red (1) sectors (Figure 2A, supplemental Figure S5). Shoot and root samples from an independent set of sand culture-grown plants were investigated for this purpose. The plants were either supplied with phosphate (+P) or were P-limited. P-limited plants were also re-supplied with phosphate for 30 min or 3 hours before harvesting. RT-qPCR demonstrated that virtually all thirty tested gene transcripts again displayed a strong P-response (supplemental Figure S5A, B). Re-addition of phosphate to P-limited plants only slightly affected the abundance of these transcripts within 30 min but led to a marked reversion for most of the gene transcripts within three hours, indicating that these PRGTs probably directly respond to a change in P status rather than an indirect and delayed change in metabolism or development. More detailed comparison of the RNA-Seq and RT-qPCR data of these 30 gene transcripts also revealed good quantitative correlation (supplemental Figure S5C, D), including the very strong induction of *AT2G34202*/*miR399d* and *INDUCED BY PHOSPHATE STARVATION1* (*AT3G09922*/*IPS1*).

### Sand-grown, P-limited *Arabidopsis* plants display many of the familiar transcriptional P-limitation responses known from plate- or liquid culture-grown younger seedlings

In the present study, we investigated 19-day old *Arabidopsis thaliana* Col-0 plants grown in a sand/perlite mix in phytotrons (Figure 1 cf. materials and methods). In terms of growth conditions and age, there is a considerable difference between the plant materials used in this study and the plate- or liquid culture-grown younger seedling materials used in previous ATH1 P-limitation studies (Misson et al., 2005; Morcuende et al., 2007; Bustos et al., 2010). Nonetheless, many previously identified PRGTs were also identified here. For example, of the 100 most strongly P-limitation induced gene transcripts detected in shoots of plate-grown seedlings by Bustos et al. (2010), 62, 24 and 12 were present in the green, orange and brown sectors, respectively (Figure 2A). Consequently, the major (hallmark) transcriptional responses to P-limitation were also identified in our study (supplemental Table S1). These hallmark transcriptional responses include induction of gene transcripts for (i) uptake and transport of Pi and other inorganic ions (*Pht1*.*3/1*.*4/1*.*5/1*.*8/1*.*9, Pht3*.*2, PHO1*;*H1, SULTR1*.*3/3*.*4, KUP6/8/10/11, GPT2, NRT1*.*8*), (ii) Pi salvage systems (12 purple acid phosphatase gene transcripts, phospholipid degradation genes *GDPD1/2/6, PEPC1, PLDZ2*), (iii) alternative metabolic pathways that lower P requirements (*AT1G73010, AT3G53620, AT4G01480, AT2G46860*) (iv) sulfo- and glycolipid synthesis (*SQD1/2, MGD2/3*), (v) redirection of carbon metabolism (*PPCK1/2, PPC3, GWD3, BAM5, APL3/4, SPS1/4*), (vi) phytohormone synthesis/response pathways (*ARR16, ERF70, GCL1*), and (vii) signaling (*SPX1/2, HRS1, HHO, PAP2/MYB90, NF-YA2/7, CYCP4*.*2, S6K2*) (supplemental Table S1).

### Functional overview of novel P-status responsive gene transcripts detected in sand-grown, P-limited *Arabidopsis* plants

To obtain a broad functional overview of the novel PRGTs in the red and pink sectors (Figure 2A), their *Arabidopsis* Gene Identifiers (AGIs) were subjected to a Gene Ontology (GO) analysis in TAIR (Figure 2C). Compared to the 229 GTs that were found to be strongly P-limitation induced in the present RNA-seq study and in the Bustos et al., 2010 ATH1 study (green sector), the 283 GTs in the red sector (identified from this study, Figure 2A) displayed a notably increased frequency of GO terms “transcription factor” (7.0%), “DNA/RNA binding” (10.2%) and “unknown molecular function” (12.8%), while the terms “hydrolase activity”, “transferase activity” and “transporter activity” were reduced. The abundance of GO term ““unknown molecular function” further increased (to 33.2%) in the set of 259 PRGTs in the pink sector (Figure 2A, novel transcript identified from this study without ATH1 probe set on ATH1 array), while the terms “hydrolase activity” (8.2%) and “transferase activity” (6.8%), further declined.

Next, individual inspection of the PRGTs in the pink and red sectors (Figure 2A) was conducted and led to several observations. The pink sector contained several previously identified PRGTs that are not represented by probe sets on ATH1 gene chips. These include phosphate transporter transcripts *Pht1*.*1* and *Pht1*.*7*, long non-coding RNAs *IPS1, IPS2/AT4* and *TAS4*, primary transcripts of microRNAs including *miR169, miR2111, miR827* and *miR399* (Pant et al., 2009), and purple acid phosphatase transcripts *PAP11* and *PAP19* (Morcuende et al., 2007). This observation further validates the P-limited status of the investigated plant materials and the usefulness of the produced RNA-seq data.

Several PRGTs in the red and pink sectors extended our understanding of transcriptional plant responses to P-limitation by increasing the number of P-responsive members in several gene families. This applies not only to *PHT1* or *PAP* transcripts mentioned above, but also to *PHOSPHOENOLPYRUVATE CARBOXYLASE* (*AT1G68750/PPC4*), *GDPD* (*GDPD3*) or *PHOSPHATE 1* (*PHO1*;*H10, PHO1*) family members, for example (Figure 2A, Table 1). Furthermore, PRGTs were identified in the pink, red and blue sectors (Table 1) that encode additional activities facilitating metabolic adaptations that occur during P-limitation and help to acclimatize plants to P deficiency through enabling Pi recycling, mobilization and scavenging, or decreasing metabolic P requirements (Plaxton and Tran, 2011). Examples include GTs encoding (i) activities that bypass metabolic pathways, such as cytosolic glycolysis (*ALDEHYDE DEHYDROGENASE 11A3/ALDH11A3/AT2G24270, MATERNAL EFFECT EMBRYO ARREST 51/MEE51/AT4G04040, PPDK, PYRUVATE ORTHOPHOSPHATE DIKINASE/AT4G15530, UDP-GLUCOSE PYROPHOSPHORYLASE 1/UGP1/AT3G03250*) and mitochondrial electron transport (*NADP-MALIC ENZYME 1/NADP-ME1/AT2G19900, ALTERNATIVE OXIDASE 1D/AOX1D/AT1G32350, AT3G33103*), and thus reduce or negate the dependence on phosphate or adenylates, (ii) activities that scavenge and recycle Pi from intra- and extracellular organic phosphorus (P) compounds (e.g. *AT1G52470, VEGETATIVE STORAGE PROTEIN 2/VSP2/AT5G24770*), (iii) intracellular (vacuolar) and secreted purple acid phosphatases (e.g. *PAP11* and *PAP19*), (iv) activities for synthesis and excretion of organic acids for soil acidification (e.g. *AT1G68750/PPC4, AT1G53310/PPC1, AT2G42600/PPC2, AT2G19900/NADP-ME1*), and (v) activities involved in breakdown of membrane phospholipids and replacement by sulfolipids and galactolipids (e.g. *AT3G03530*/*NPC4, AT2G01180, AT3G11670*/*DGD1*).

**Table 1.**
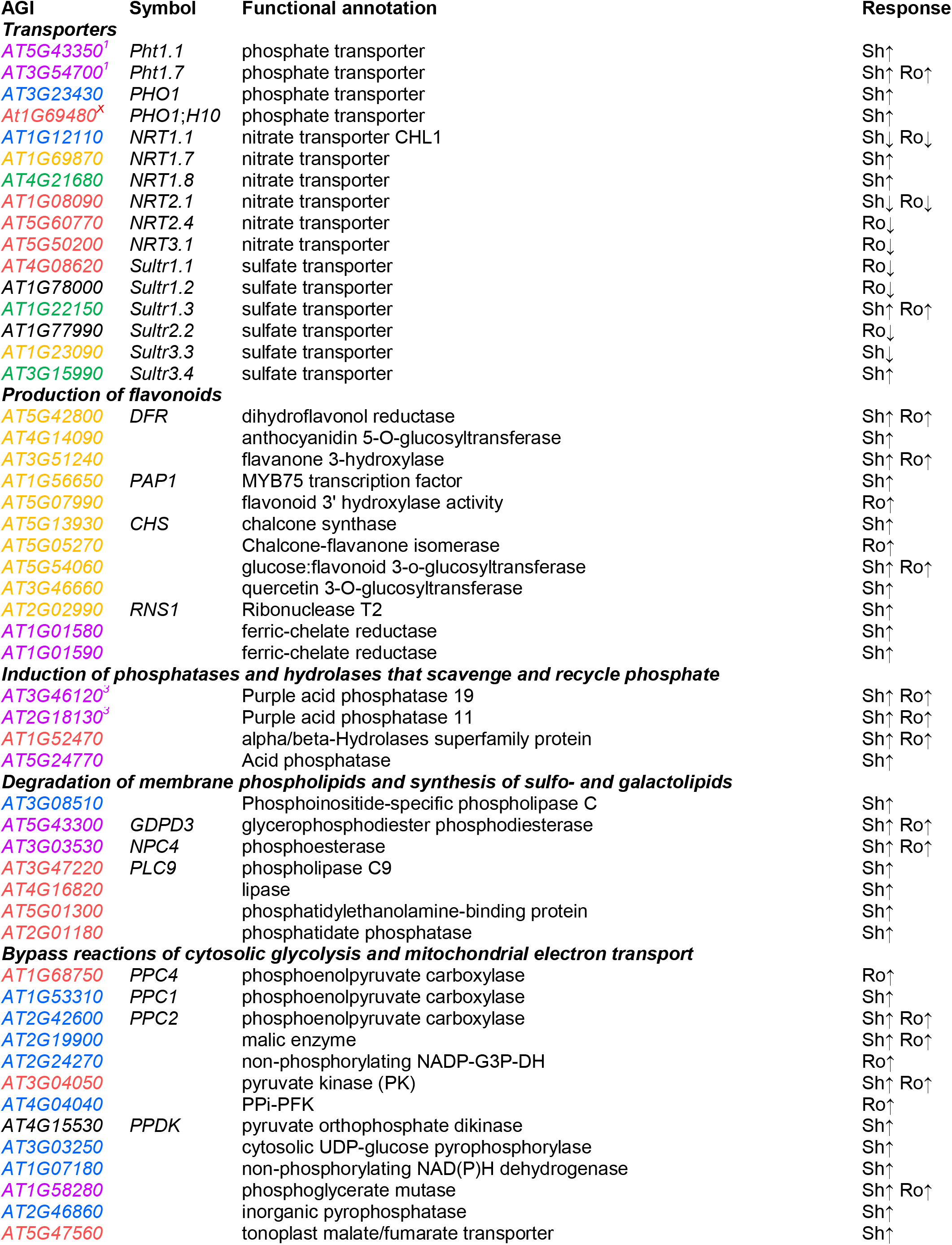
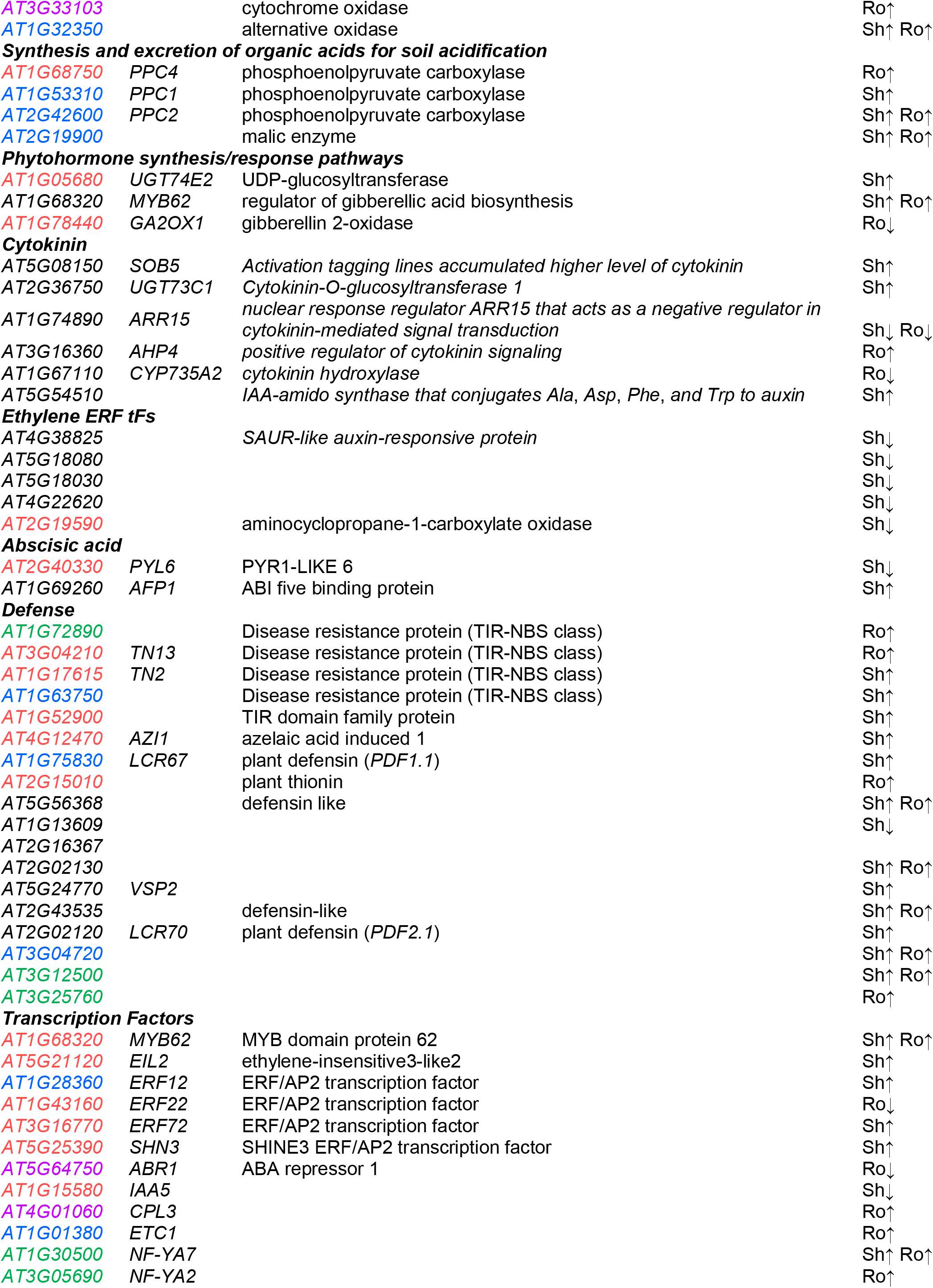
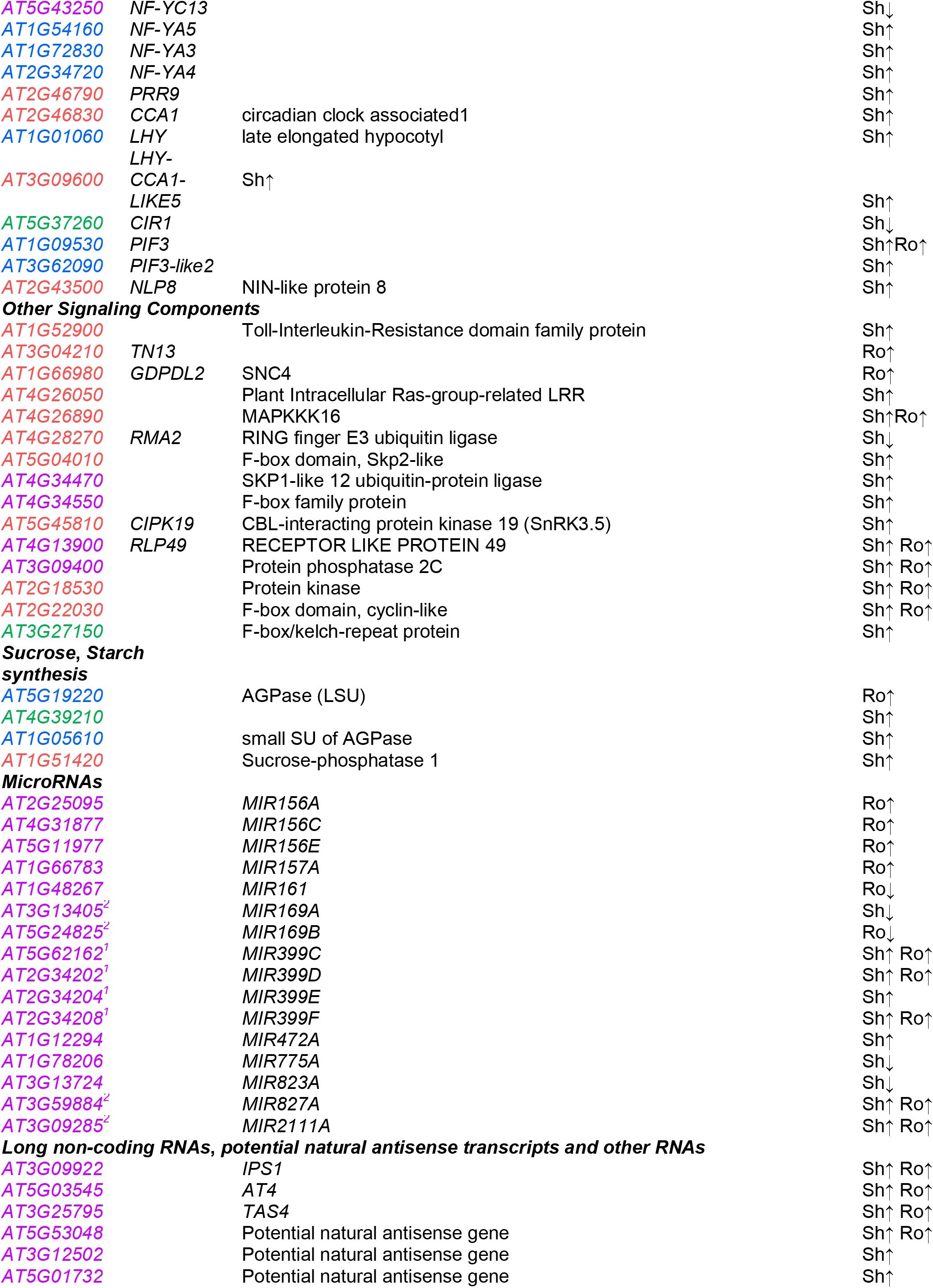
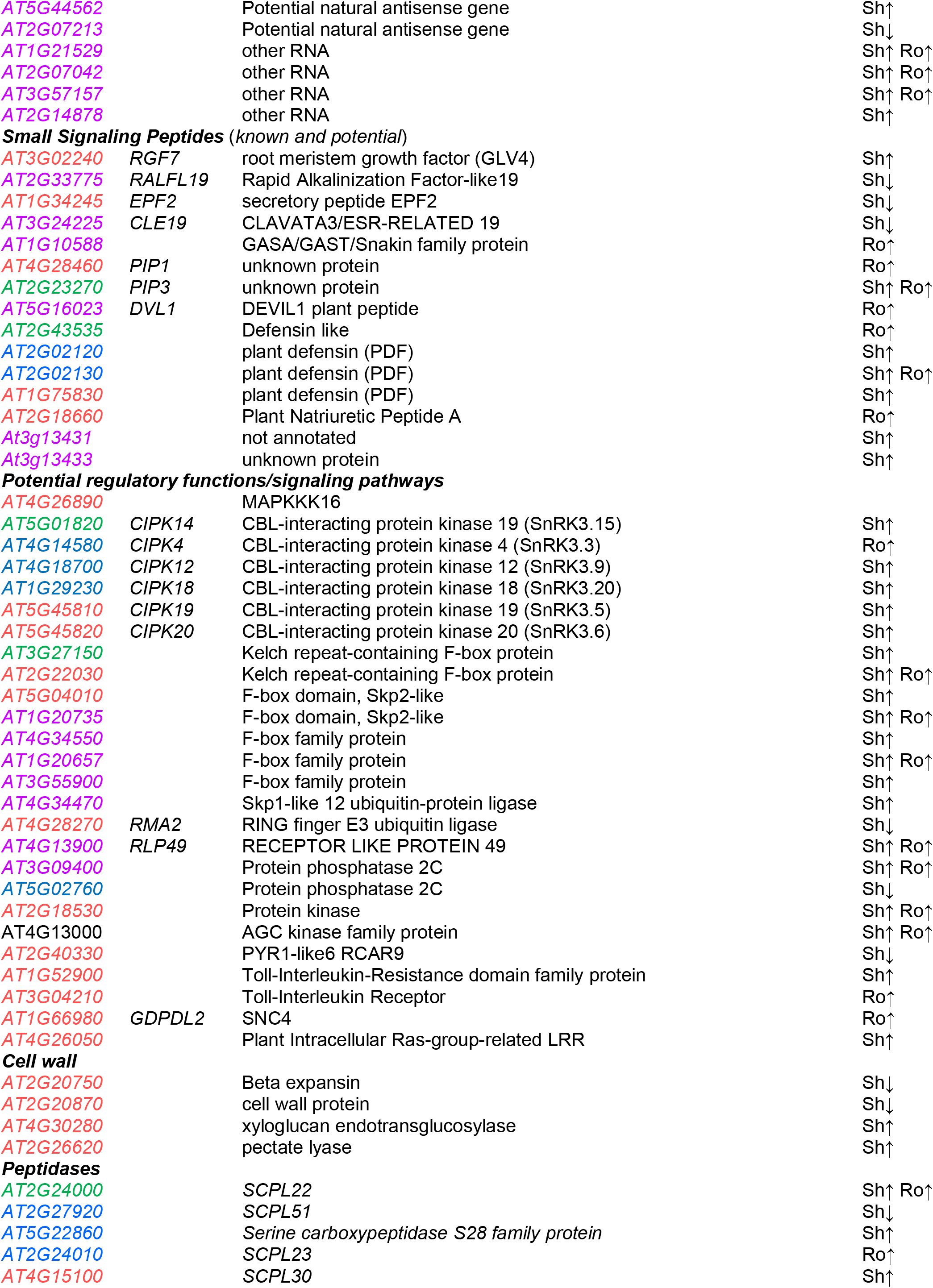
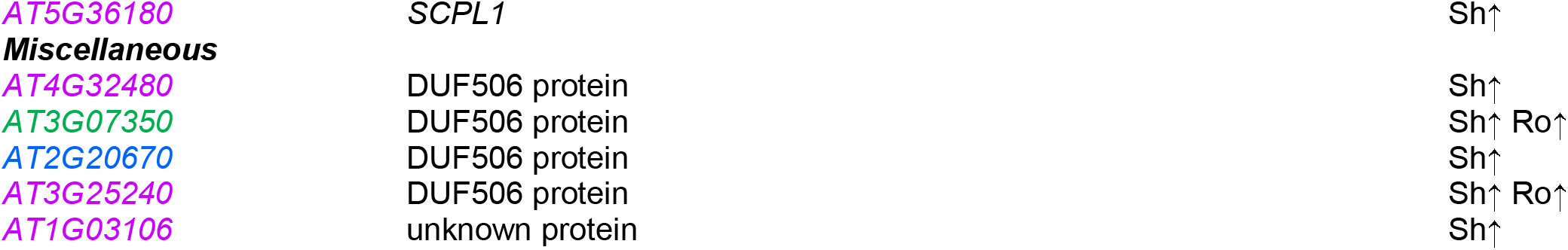
Overview of novel and previously identified P-status responsive gene transcripts (PRGTs). As described in Figure 2, PRGTs are shown in pink (≥5-fold changed in RNA-seq and without probe set on ATH1 study), blue (≥5-fold changed in RNA-seq and 2-5-fold changed in ATH1 study), red (≥5-fold changed in RNA-seq without support from ATH1 study), yellow (PRGTs of yet unannotated genes), and green (PRGTs ≥5-fold changed in RNA-seq and ATH1 study. Transcripts labeled with superscripts were previously reported as P-status responsive in the respective reference: (1) Bari et al. (2006), (2) Pant et al. (2009), (3) Morcuende et al. (2007).

### Novel PRGTs encoding transporters

Based on ATH1 Affymetrix gene chip studies, P-limitation is known to affect expression of ion transporters, most notably to induce *PHT1* phosphate transporters, but also potassium and sulfur transporters, indicating the profound effect of P-limitation on ion homeostasis. PRGTs in this study with GO annotation “transporter activity” in the blue, red and pink sectors (Figure 2A) comprised phosphate transporters transcripts *PHT1*;*7* and *PHT1*;*1* (see above; Figure 2A), as well as *PHO1* (*AT3G23430*; see blue sector) and its homolog *PHO1*.*H10*; (*At1G69480*). More surprisingly, four PRGTs from the blue and red sectors, encoding high-affinity nitrate transporters, i.e. *NRT1*.*1* (*AT1G12110*), *NRT2*.*1* (*AT1G08090*), *NRT2*.*4* (*AT5G60770*) and *NRT3*.*1* (*AT5G50200*) were strongly repressed in roots, suggesting that P-limitation leads to a decrease of nitrate uptake. *NRT1*.*1* and *NRT2*.*1* also showed repression in shoots, whereas *NRT1*.*7* (*AT1G69870*) and *NRT1*.*8* (*AT4G21680*) were induced. Repression in P-limited roots was also found for *AT4G08620, AT1G77990* and *AT1G78000* encoding sulfate transporters 1.1, 2.2 and 1.2, respectively, whereas induction of sulfate transporters 1.3 (*AT1G22150*) and 3.4 (*AT3G15990*) is known from ATH1 Affymetrix gene chip studies and was confirmed here (see above). In addition to these changes, the RNA-seq analyses also revealed novel PRGTs encoding other cation/anion, peptide, sugar, ABC, MtN21/UMAMIT and MATE transporters (Table 2). Particularly noteworthy are the 10 P-status responsive transcripts encoding members of the MtN21/UMAMIT family of transporters. Three of these (*AT4G08300, AT4G28040, AT5G50800*/*SWEET13*) were already known from ATH1 studies to be P-responsive (), while another seven, five induced (*AT1G21890, AT2G39510, AT4G15540, AT5G07050, AT5G13170*/*SWEET15*/*SAG29*) and two repressed (*AT1G60050, AT1G70260*), were identified from this study (Table 2).

**Table 2.** PRGTs encoding transporters. PRGTs are shown in green (≥5-fold changed in present RNA-seq and ATH1 study), blue (≥5-fold changed in RNA-seq and 2-5-fold in ATH1 study), red (≥5-fold changed in RNA-seq but not changed in ATH1 study) and pink (≥5-fold changed in RNA-seq and no probes set present on ATH1 gene chips).

| Phosphate | Nitrate | Sulfate | K Na | Ca Cu Fe | Sugars | Amino acids | Peptides | MtN21<br>UMAMIT | ABC | MATE |  |
| --- | --- | --- | --- | --- | --- | --- | --- | --- | --- | --- | --- |
| Pht1.2, Pht1.3<br>Pht1.4, Pht1.5<br>Pht1.8, Pht1.9<br>GPT2<br>G3PP1 | NRT1.8 | SULTR1.3<br>SULTR3.4 | KUP6<br>KUP8<br>KUP10<br>KUP11<br>NHX2 | CAX3<br>COPT2 | ESL1<br>SWEET13 | CAT3<br>At1g08230 |  | UMAMIT17<br>UMAMIT33 | ABCC10<br>ABCG22 | At1g33110 | BOR5 PUP4<br>YSL8 AMT1.2<br>OCT1 OCT4<br>DTX1 AHB2<br>MRP14 DIC3 |
| PHO1<br>G3PP3 | NRT1.1<br>NRT3.1 | SULTR2.2 | SKOR<br>KEA2<br>CHX17 | VTI2 (ABG4) | STP7<br>SWEET15 | BAC2<br>At5g02180 | NPF6.4<br>OPT3 |  | ABCG1<br>ABCG34 | ABS4<br>At1g66700<br>At1g73700<br>At2g04050<br>At2g04100 | At1g72700<br>PILS3 TIP1.3<br>ALA2 |
|  | NRT2.1<br>NRT2.4 | SULTR1.1<br>SULTR1.2 | HAK5<br>SLAH1 | CHX21<br>At3g08040<br>ZIP12 | SUC7 | GLR2.3<br>GLR2.9<br>At2g41190 |  | UMAMIT9<br>UMAMIT14<br>UMAMIT19 | ABCB15<br>ABCG27<br>ABCC9<br>ABCE3<br>ABCB7 | At2g04070<br>FRD3<br>DTX50 | TDT |
| Pht1.1 Pht1.7<br>At4g08878 |  |  | SLAH4 |  |  | GLR2.2<br>At5g16740 |  | UMAMIT35<br>UMAMIT36<br>UMAMIT38<br>UMAMIT40 | ABCB11<br>ABCC3 | At2g04090<br>BIGE1A | ENT3<br>MSL4<br>At1g04570<br>At2g23790 |

### Novel PRGTs that encode transcription factors

PRGTs annotated as “encoding transcription factors” were analyzed in more detail, due to their importance and because of their abundance (33, i.e. 11.7%) in the red sector (Figure 2C and supplemental Table S2), suggesting that the sequencing depth and sensitivity of RNA-seq was high enough to detect gene transcripts with low average expression (supplemental Figure S3), such as those encoding transcription factors previously missed using ATH1 arrays. In the green, blue and pink sectors (figure 2A) 11 (4.8%), 19 (9.6%) and 11 (4.2%), respectively, PRGTs have the GO annotation “transcription factor” yielding a total of 74 P-responsive transcription factor gene transcripts (TFGTs) identified by RNA-Seq. Sashimi plots of eight novel P-limitation induced and two repressed TFGTs, four (*AT1G18860, AT1G49900, AT1G60280, AT2G46790*) from the red and four (*AT1G27045, AT3G18010, AT3G28857, AT5G55690*) from the pink sector (Figure 2A) are shown in Figure 3. While the RNA-seq results confirmed the existing annotations of genes like WUSCHEL-related homeobox *AT3G18010* and *AT2G46790*/*PRR9* (Figure 3B, G), the Sashimi plots for *AT5G55690* (Figure 3A), the highly induced MADS box TF gene transcript *AGL47*, previously identified as P-responsive by RT-qPCR (Morcuende et al., 2007), and *AT1G18860* (Figure 3E), a WRKY TF, both revealed an additional unannotated exon. Sashimi plot for *AT1G49900* (C2H2 zinc finger TF; Figure 3D) suggests that the current annotation encompasses two distinct PRGTs. *AT1G27045*/*HB54* (Figure 3F) and *AT1G60280/NAC23* (Figure 3H) are two examples of TFGTs with very low expression, yet their P-response was clearly identified using RNA-seq (supplemental Table S1).

**Figure 3.**
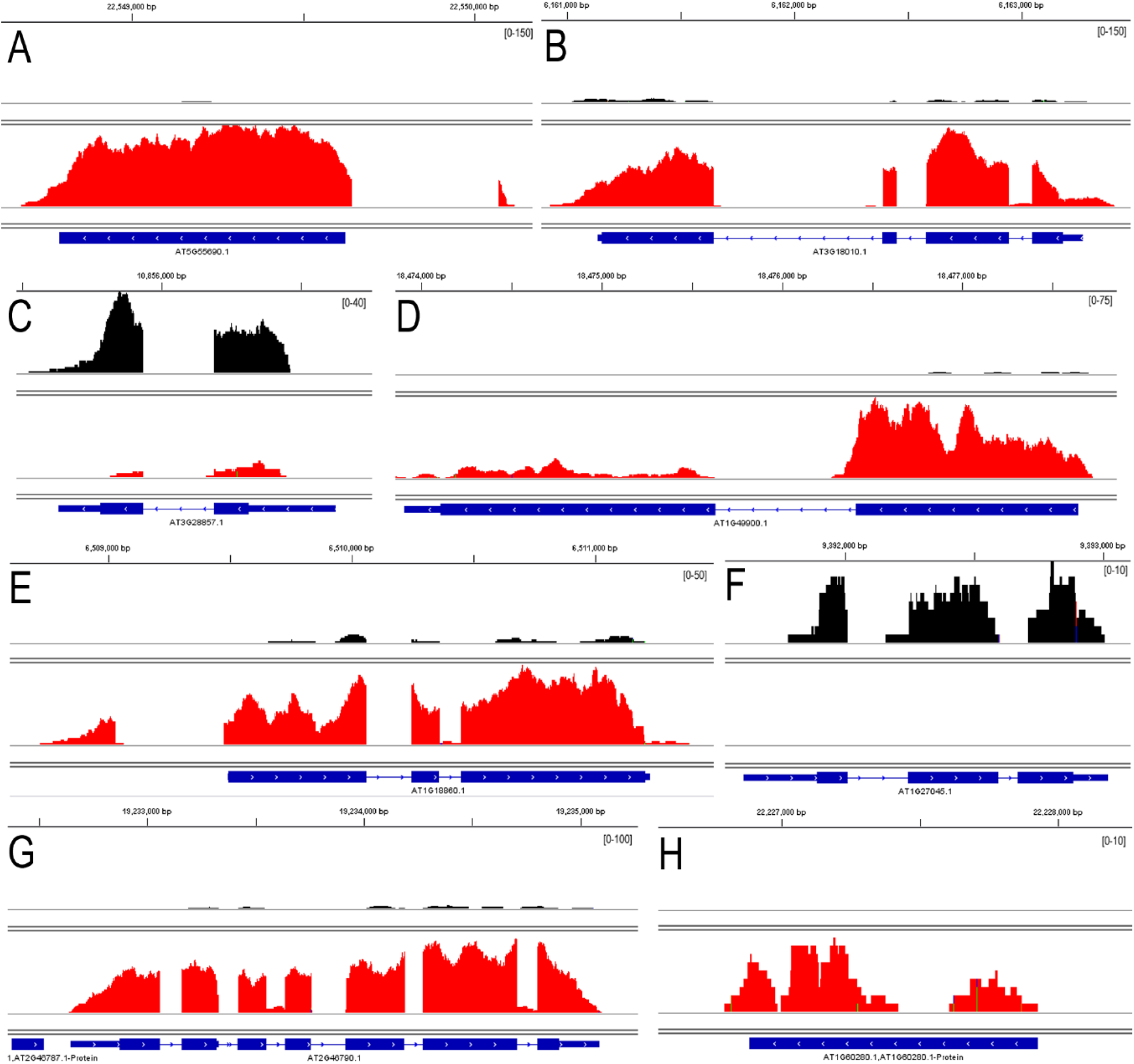
Sashimi plots for selected novel P-status responsive transcription factor gene transcripts. Panels (A, *AT5G55690*; B, *AT3G18010*; C, *AT3G28857*; D, *AT1G49900*; E, *AT1G18860*; F, *AT1G27045*; G, *AT2G46790*; H, *AT1G60280*) were produced with IGV and BAM-files originating from RNA sequencing of P-limited (red) and P-sufficient (black) shoot samples. Scaling of FPKM read numbers is the same for the plots of the same transcript as indicated in parentheses (upper right corner of each panel).

Among the 63 P-responsive TFGTs from the blue, red and pink sectors (Table 1, Figure 2A), there are a few with a remarkably strong P-response (e.g., *AT1G49900* and *AT5G55690*) and some for which biological functions are known to some extent. Several TFGTs provide links between P-limitation and hormone signaling, such as *AT1G68320*/*MYB62, AT5G21120*/*EIL2*, ethylene response factors (*AT1G28360*/*ERF12, AT1G43160*/*ERF22, AT3G16770*/*ERF72, AT5G25390*/*SHN3*), *AT5G64750* encoding ABA REPRESSOR1, or *AT1G15580*/*IAA5*, while *AT4G01060*/*CPL3* and *AT1G01380*/*ETC1* link P-limitation to root hair development. Besides two TFGTs encoding nuclear factor Y subunits (*AT1G30500, AT3G05690*) from the green sector, four additional ones (*AT5G43250, AT1G54160, AT1G72830* and *AT2G34720*) were present in the blue and pink sectors. Furthermore, four TFGTs (*AT2G46790*/*PRR9, AT2G46830*/*CCA1, AT1G01060*/*LHY, AT3G09600*/*LHY-CCA1-LIKE5* involved in clock and circadian function were considerably induced during P-limitation (Figure 3G). In addition, *AT1G09530*/*PIF3* which interacts with promoter elements of *LHY* and *CCA1* and with photoreceptors PhyA and PhyB, and *AT3G62090*/*PIF3-like2* which physically associates with TOC1/PRR1, a pseudo response regulator involved in the generation of circadian rhythms, were induced. It thus appears that P-limitation can affect clock and circadian functions through several of the known, central regulators.

### Novel PRGTs involved in signaling and regulatory processes

In addition to the above mentioned PRGTs, we sought for other novel potential regulatory gene transcripts that respond to P availability. These included gene transcripts encoding (i) regulatory RNAs (microRNAs, long-noncoding RNAs), (ii) proteins with functions in signaling, including receptor kinases, small secreted peptides, and MAP kinases, and (iii) proteins with predicted functions in post-translational protein modification (kinases and phosphatases) and protein degradation for example F-box proteins and E3 ligases (Table 1).

Primary transcripts of sixteen microRNAs were detected among the 5-fold changed gene transcripts (supplemental Table S1, Table 1). While induction of *miR399b-f, miR827, miR2111a* transcripts was very strong and expected (Pant et al., 2009), the induction in roots of four primary transcripts encoding *miR156a, c, e* and *miR157a* (all targeting SPL transcription factors) and repression of two transcripts encoding miR169a/b (targeting NF-YA HAP-type transcription factors) in shoots and roots, respectively, was less pronounced and unreported. Repression was also observed for the primary transcripts encoding *miR161, miR823a* and *miR775a*, while the one for *miR472a* was induced. Besides microRNAs, 11 long non-coding RNAs (lncRNAs) were identified among the genes in the pink sector (supplemental Table S1, Table 1), including the known *IPS1, AT4*/*IPS2* and *TAS4*, four genes annotated as “potential natural antisense RNA” (*AT3G12502, AT5G01732, AT5G44562, AT5G53048*) and another four annotated as “other RNA” (*AT1G21529, AT3G57157, AT2G14878, AT1G43765*).

Gene transcripts encoding small secreted peptides (SSPs), known to play a role in many signaling pathways, were (and continue to be) poorly annotated and thus hardly represented on ATH1 gene chips. During recent years the importance of SSPs emerged especially with regard to their crucial developmental functions, and more recently also with regard to their role in plant nutrition (Tabata et al., 2014; de Bang et al., 2017; Ohkubo et al., 2017). The roles of SSPs during P-limitation are yet uncharacterized and P-status responsive *SSP* gene transcripts are thus far almost undescribed. We detected fifteen *SSP* transcripts among the 1,074 PRGTs (supplemental Table S1; Table 1), including those encoding root meristem growth factor RGF7/GLV4, rapid alkalinization factor RALFL19, EPF2, PIP1 and PIP3, CLE19, DVL1 and three plant defensins. Moreover at least two PRGTs (*AT1G08165, AT3G13433*) encoding potential novel peptides were identified among the 1,074 PRGTs. Inspection of PRGTs with responses <5-fold (Figure 1) yielded a total of 38 annotated *SSP* transcripts with at least 2-fold changed abundance in shoots and roots during P-limitation. It will be important to investigate how P-status responsive SSPs, either produced *in planta* or synthetically, affect the plant P-limitation response including root development, and what their cognate peptide receptors (usually LRR-RLKs) are.

Over 20 additional PGRTs with potential regulatory functions were identified in this study as shown in the blue, red and pink sectors (Figure 2, supplemental Table S1; Table 1). For example, *AT4G26890*, encoding MAP3K16, was ∼9-fold induced in shoots, and, in addition to MAP3K18 and 19 (*AT1G05100, AT5G67080*) which are present in the green sector, represents the third MAP3K responsive to P-status. The transcript encoding calcineurin B-like-interacting serine/threonine protein kinase 14 (*CIPK14, AT5G01820*) was known from ATH1 studies to be P-limitation induced in shoots, and this induction was confirmed here. More interestingly, five additional CIPK transcripts (4, 12, 18, 19 and 20; Table 1) were identified in this study to be induced by P-limitation in shoot (CIPK12, 18, 19, 20) or root (CIPK4), thus suggesting that several signaling pathways involving CIPKs calcium sensors are modulated by P-status. Furthermore, at least six novel genes encoding P-responsive F-box proteins, a Skp1-like ubiquitin protein ligase, a RING-finger E3 ubiquitin ligase, two protein phosphatases 2C, as well as PYR1-like 6, a regulatory component of the abscisic acid receptor, were found to respond to P-status at the transcript level (supplemental Table S1; Table 1).

### Novel PRGTs encoding cell wall, peptidases and domain of unknown function proteins

Over thirteen novel PRGTs related to cell wall, peptidases, domain of unknown function (DUF506) and unknown proteins were identified in this study (supplemental Table S1; Table 1). This includes four gene transcripts in red sector such as *AT2G20750/Beta expansin* and *AT2G20870/cell wall protein* that were repressed in shoots and *AT4G30280/xyloglucan endotransglucosylase* and *AT2G26620/pectate lyase* that were induced in shoots during P-limitation. Differential regulation of these cell wall related genes fits well with fine-tuning of cell wall formation during P-limitation. Five additional PRGTs encoding peptidases (*AT5G36180*/*SCPL1, AT2G24010*/*SCPL23, AT4G15100/SCPL30, AT2G27920/SCPL51, AT5G22860/Serine carboxypeptidase S28 family*) were identified in this study and included in blue, red and pink sectors (supplemental Table S1; Table 1). P-limitation induction of previously known *AT2G24000*/*SCPL22* was confirmed here. Other novel PRGTs identified by this study include DUF506s or unknown protein encoding gene transcripts such as *AT3G25240, AT4G32480, AT2G20670*, and *AT1G03106* as shown in blue and pink sectors (supplemental Table S1; Table 1, Figure 2A). *AT3G07350/DUF506* was known from ATH1 studies to be P-limitation induced in shoots and roots, and this induction was confirmed here.

### Unannotated transcriptional units, long non-coding RNAs, and a novel P-responsive microRNA

Unannotated transcripts (106 in total) with >5-fold change in abundance during P-limitation were identified (Figure 2A; yellow sector). Sequences of these unannotated PRGTs were extracted and further examined. The median length was 522 nt with the longest reaching ∼1700 nt and the shortest ∼150 nt (Figure 4A). More than 80% of the unannotated PRGTs have no intron, ∼13% have one intron, and one of the transcripts (MSTRG.19564) has five introns (Figure 4A insert). Several unannotated transcripts (Table 3, supplemental Figure S6 and S7) such as *MSTRG*.*8269* (Figure 4B), *MSTRG*.*13996* (Figure 4C), *MSTRG*.*11040* (Figure 4D), *MSTRG*.*19189* (Figure 2B), or *XLOC_005412* and *XLOC_001234*) are very strongly induced and thus highly expressed during P-limitation (Figure 1B; supplemental Table S1). The unannotated transcripts have low or no coding potential, i.e. only small or no open reading frames are discernible, thus classifying them, *a priori*, as long non-coding RNAs (lncRNAs). A comparison with the set of ∼1200 lncRNAs suggested by Yuan et al. (2016) revealed that the genome coordinates of 21 unannotated PRGTs overlapped with the coordinates of one of these lncRNAs, and only five unannotated PRGTs (*MSTRG*.*14089, MSTRG*.*19189, MSTRG*.*19313, MSTRG*.*21504, XLOC_005412*) were found to be P-responsive (≥5-fold) by Yuan et al. (2016) (supplemental Figure S8, Supplemental Table S3 and S4), suggesting considerable differences in the P-response between both studies (see below for more discussion).

**Figure 4.**
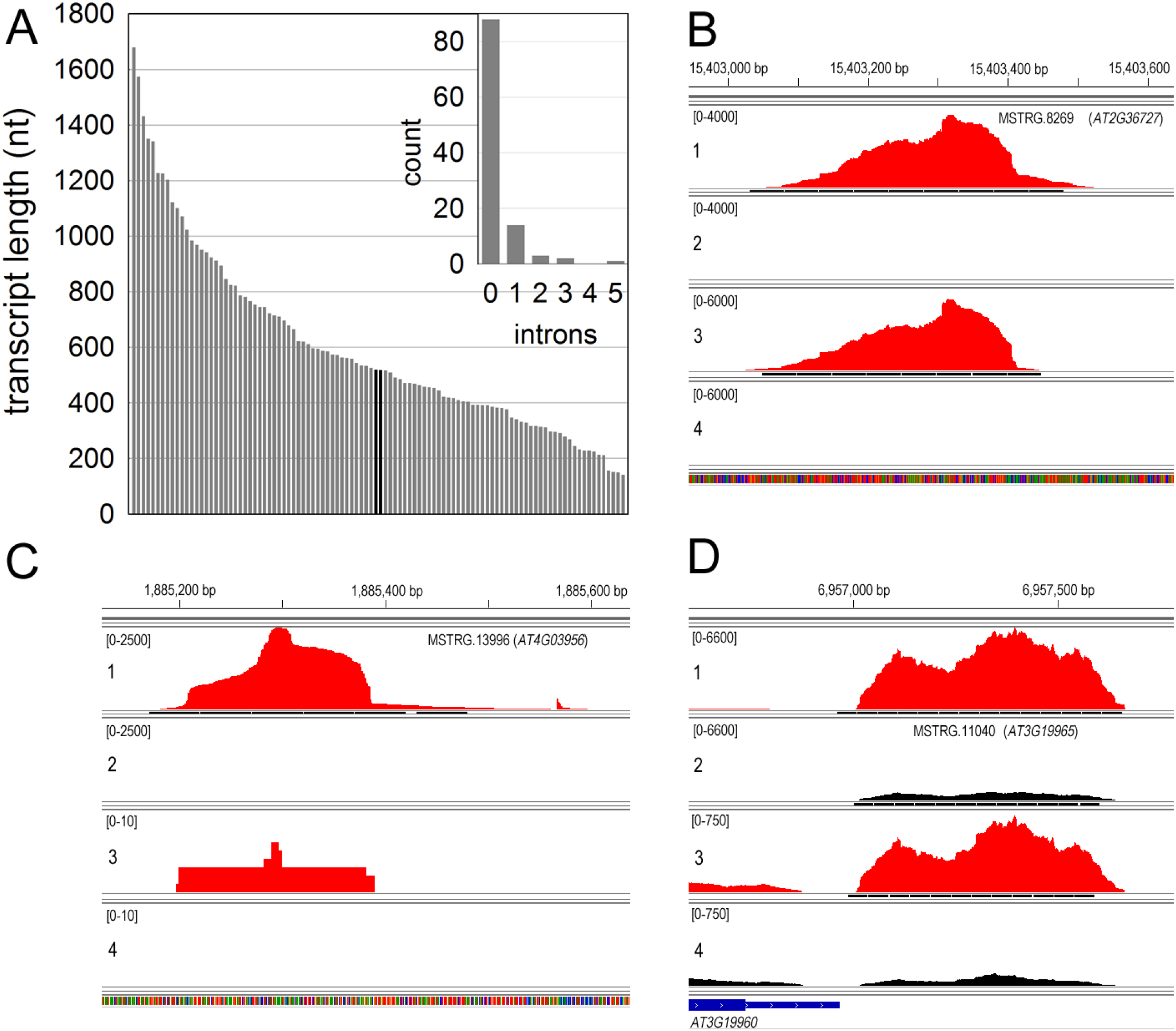
Unannotated transcripts. (A) Transcript length (nucleotides, nt) of the unannotated novel PRGTs identified in present study. Length of these transcripts range from ∼1700 nt for the longest and ∼150 nt for the shortest with median length 520 nt. Insert shows the number of transcript count (y-axis) vs. number of introns (x-axis) revealing that most of the genes have no introns. (B), (C) and (D) shows Sashimi plots for the novel P-limitation inducible gene transcripts MSTRG.8269, MSTRG.13996, and MSTRG.11040, respectively. Plots were generated using IGV and BAM-files originating from RNA sequencing of the P-limited (-P, red) and P-sufficient (+P, black) shoot samples. Scaling of FPKM read numbers for each transcript is shown in parentheses (upper left corner of each panel) in each panel (1) P-limited shoot; (2) P-sufficient shoot; (3) P-limited root and (4) P-sufficient root.

**Table 3.**
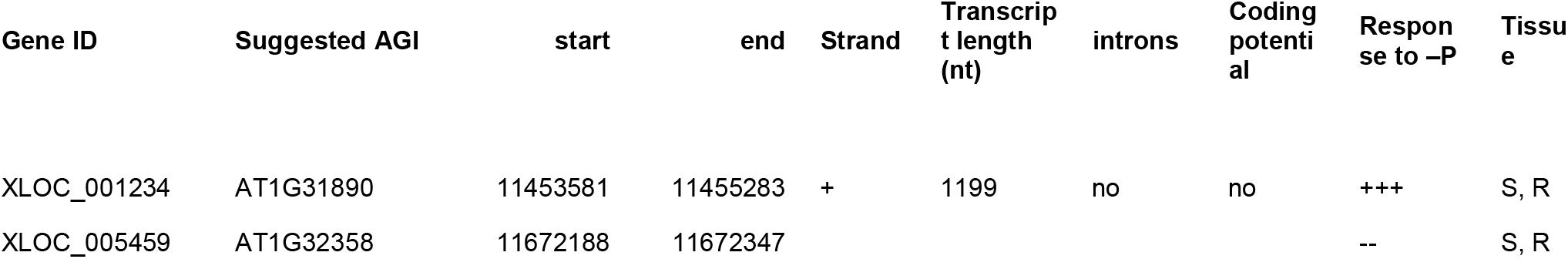
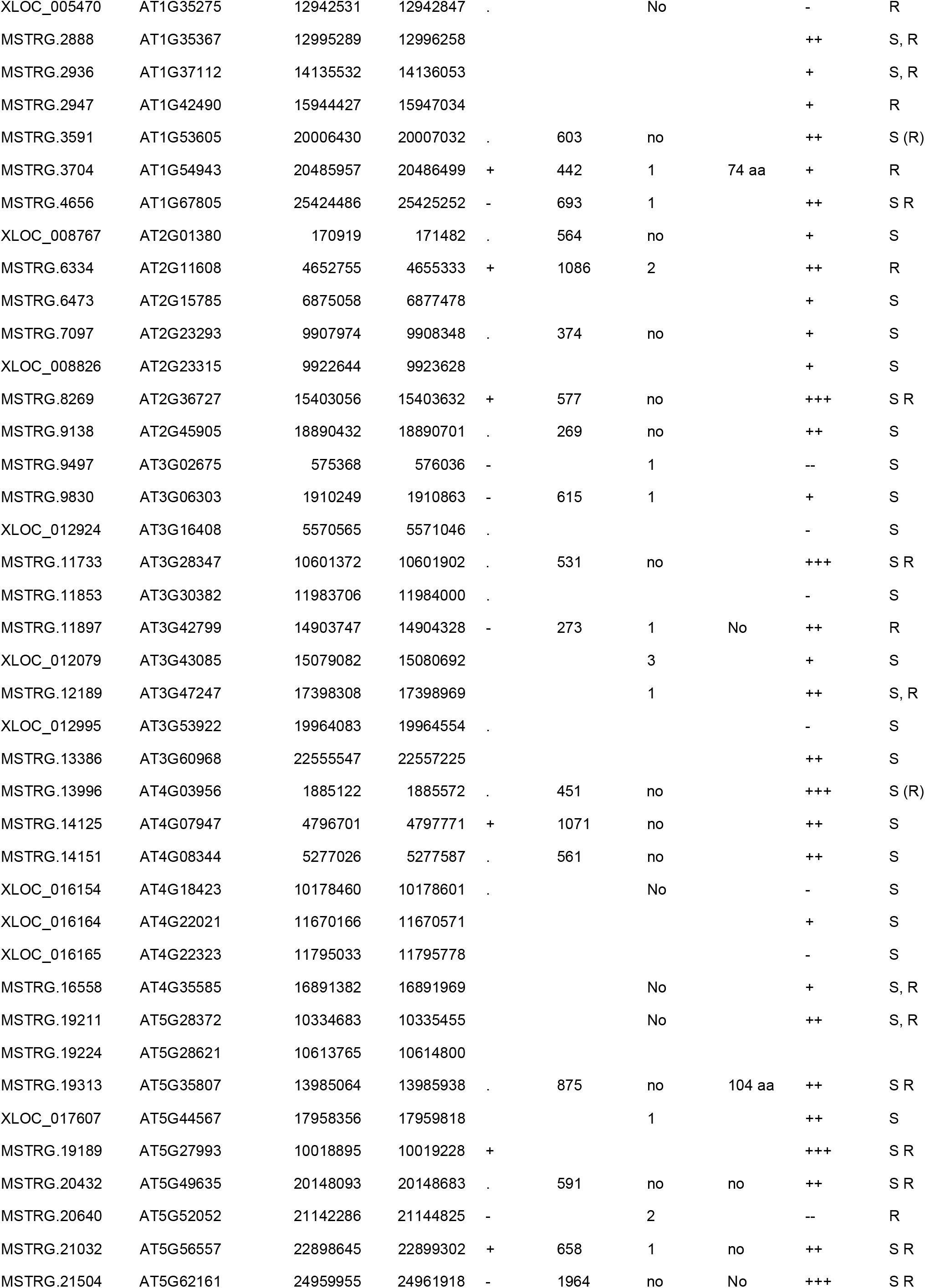
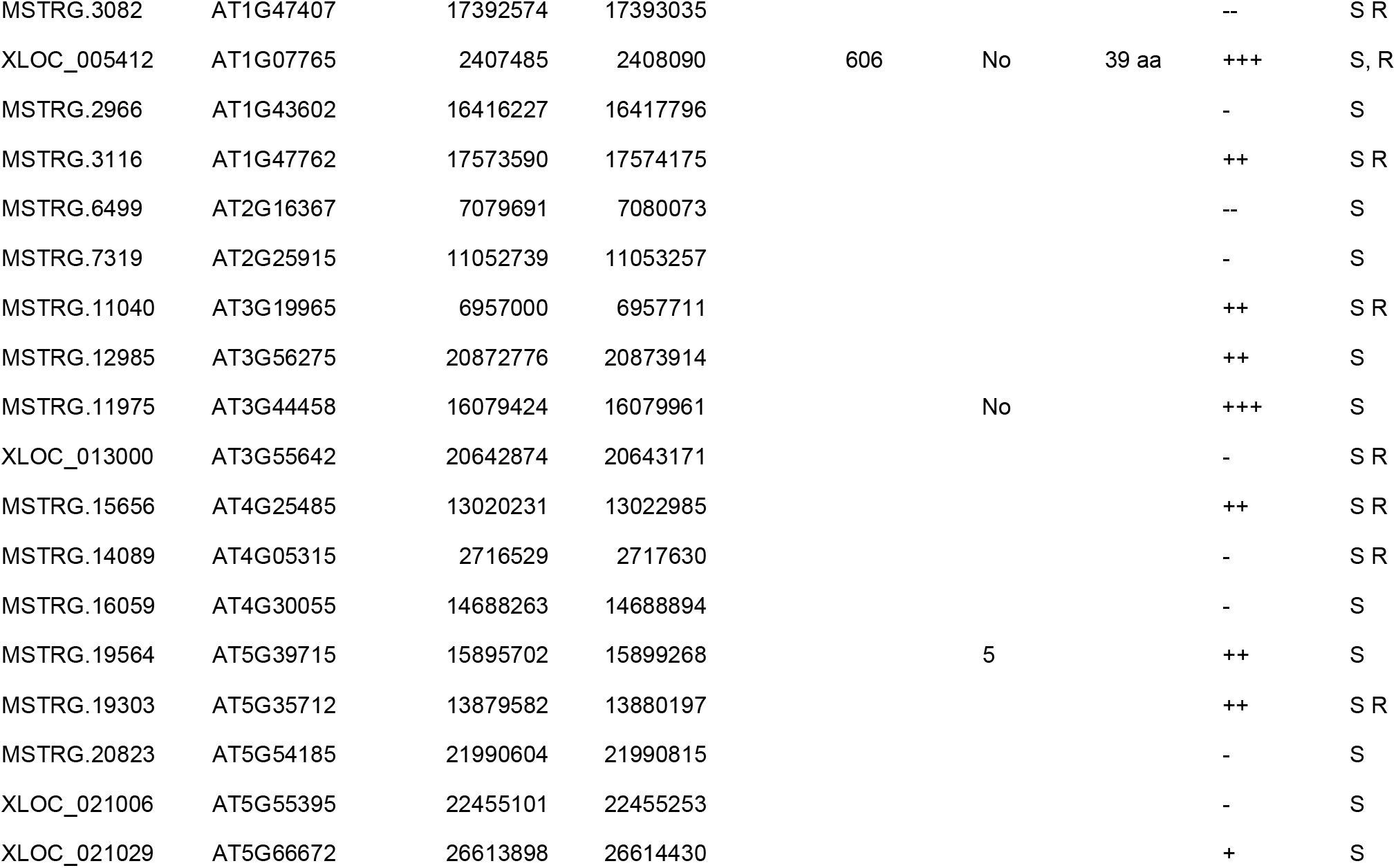
Examples of highly P-responsive, unannotated transcriptional units.

LncRNAs have various biochemical and molecular functions as target mimics (e.g. *IPS1*) (Franco-Zorrilla et al., 2007), small RNA precursors, antisense RNAs (blocking recognition of exons by the spliceosome; generating endo siRNAs) or binding partners of proteins (thus affecting protein activity, alter protein localization or providing a structural/organizational role) (Wilusz et al., 2009). Although in-depth structural analyses of the lncRNAs or their functional assignments are not within the scope of this study, the highly P-limitation induced and expressed MSTRG.19189 (*AT5G27993*) transcript was found to be produced from a genome region that can form an extended stem-loop structure (Figure 5A-C). Furthermore, a 21 nt long sequence (UCAUAAACCGUGAUUGUCCGU) present in the stem loop is highly abundant in small RNA libraries from P-limited seedlings (Pant et al., 2009) (Figure 5D), suggesting that this sequence is a yet undescribed P-limitation induced microRNA. Strong expression during P-limitation of the predicted stem-loop sequence was confirmed by RT-qPCR in three independent biological replicates (Figure 5E). Prediction of potential target genes with TargetMiner (Bandyopadhyay and Mitra, 2009) yielded three genes encoding a leucine-rich repeat protein kinase (*AT2G45340*), an F-box family protein (*AT4G12820*), and an unknown protein (*AT1G66190*) (Figure 5F).

**Figure 5.**
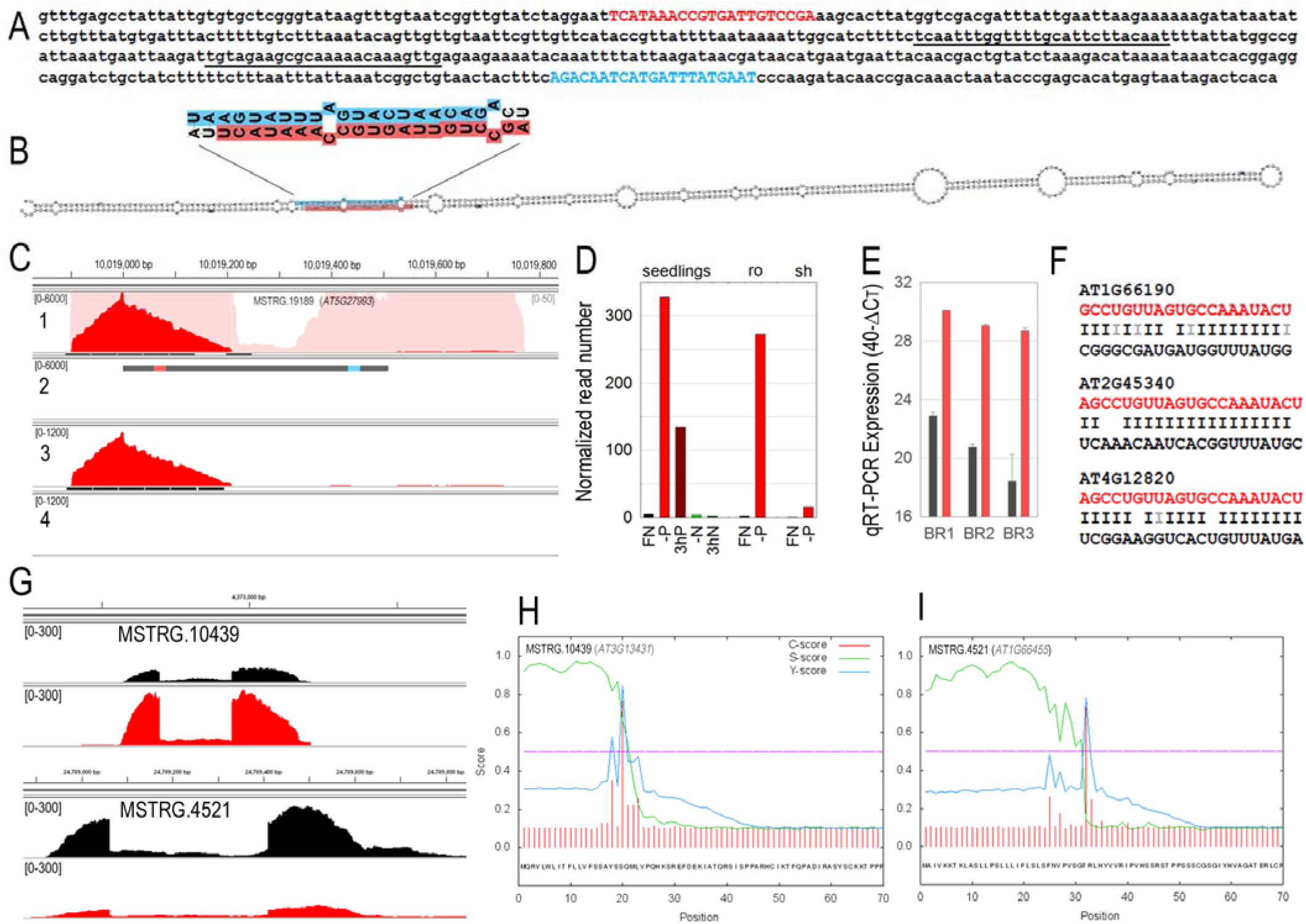
Evidence of a novel, unannotated P-starvation inducible *miRNA* gene. (A, B) Sequence and proposed folding of the microRNA stem-loop precursor. The potential mature microRNA (miR) is depicted in red, and the complementary miR* in blue. (C) Sashimi plot of RNA-seq reads covering the potential miR precursor shown in lane 2. Read integral in (1) P-limited shoot; (2) P-sufficient shoot; (3) P-limited root and (4) P-sufficient root. Numbers in brackets to the left of each lane indicate the scaling. The pale red background in lane 1 shows the read integral at 120x magnification. (D) Normalized abundance of the 21 nt microRNA sequence UCAUAAACCGUGAUUGUCCG in small RNA libraries from nutrient-replete (FN), P-limited (-P) or N-limited (-N) *Arabidopsis* seedlings, roots (ro) and shoots (sh). (E) RT-qPCR expression analysis of the potential primary microRNA transcript in three biological replicates (BR) of nutrient-replete (black bars) or P-limited (red bars) *Arabidopsis* seedlings using primer sequences as underlined in panel A. (F) Target gene prediction; Wobble base pairs are indicated by grey lines. (G) Sashimi plot of RNA-seq reads covering the potential small secreted peptides (SSPs) encoding gene transcripts MSTRG.10439 and MSTRG.4521 using IGV and BAM-files originating from RNA sequencing of P-sufficient (+P, black) and P-limited (-P; red) shoot samples. Scaling of FPKM read numbers is the same for all four plots as indicated in parentheses (upper left corner of each panel). (H) and (I) SignalP-4.1 prediction of the MSTRG.10439 and MSTRG.4521 as small secreted peptides (SSPs, Sec/SPI). Cleavage site for MSTRG.10439 is between position 19 and 20 (AYS-SQ) with probability 0.7681, and cleavage site for MSTRG.4521 is between position 31 and 32 (VSG-TR) with probability 0.8544.

The limited coding capacity of some unannotated transcripts also led us to investigate whether some of these have the potential to encode for preproproteins of small secreted peptides (SSPs) which might act as ligands of receptor like kinases in signaling pathways. One of the features of signaling SSPs is the presence of a discernible N-terminal signal peptide that targets the preproprotein to the secretory pathway that can be predicted using SignalP software. Interestingly, MSTRG.10439 and MSTRG.4521 displayed this feature (Figure 5G-I).

### Majority of the novel PRGTs are regulated by *PHR1, PHL1* and *PHO2*

Many PRGTs that are represented on the ATH1 array are known to be regulated by the MYB transcription factors *PHOSPHATE STARVATION RESPONSE1* (*PHR1*) and *PHR1-like* (*PHL1*) (Rubio et al., 2001; Bari et al., 2006; Bustos et al., 2010) and *PHOSPHATE2* (*PHO2*) (Bari et al., 2006). In order to check if a subset of PRGTs identified in the present RNA-seq study were also under the control of *PHR1, PHL1* and/or *PHO2*, RT-qPCR was performed on *phr1, phr1phl1* and *pho2* mutants (Figure 6). Out of the 30 PRGTs analyzed, 26 (87%) genes were down-regulated in *phr1phl1*, 21 (70%) genes were down-regulated in *phr1*, and 20 (67%) genes were up-regulated in *pho2* mutants in response to P. Since PHR1 is known to regulate phosphate starvation-inducible (*PSI*) genes by binding to their PHR1 binding site (P1BS) element that is present in the promoters of most *PSI* genes (Rubio et al., 2001), these 30 genes were also inspected for the presence of P1BS element in their 3000 bp upstream region. Nearly 25 out of 30 genes (83%) had PHR1 binding site in their upstream sequence and 87% of these genes were found to be down-regulated in *phr1phl1* mutants (Figure 6, supplemental table S5). These results indicate that the majority of these PRGTs act downstream of *PHR1/PHL1* and *PHO2* signaling pathway.

**Figure 6.**
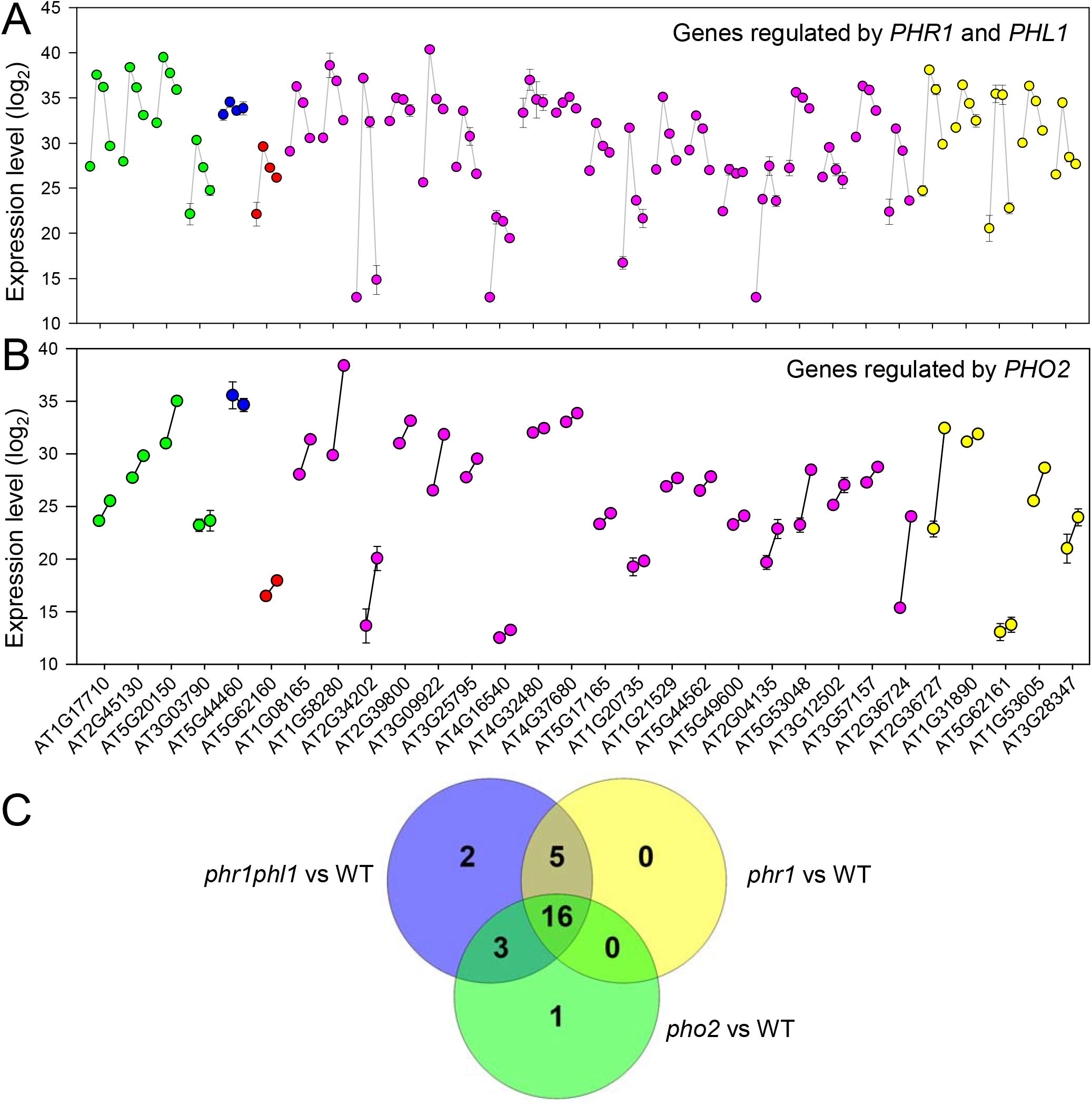
*PHR1, PHL1* and *PHO2* regulation of PRGTs identified by RNA-seq. Quantitative reverse transcription PCR (RT-qPCR) analysis of 30 PRGTs identified by RNA-seq. (A) Four data points for each gene represents its expression in P-sufficient wild-type (WT; 1^st^ data point), P-limitation WT (2^nd^ data point), P-limitation *phr1* (3^rd^ data point) and P-limitation *phr1phl1* (4^th^ data point). (B) Two data points for each gene transcript represents its expression in P-sufficient WT (1^st^ data point) and P-sufficient *pho2* (2^nd^ data point). Colors of the symbols correspond to the color coding used in Figure 2A and S5, in short, four gene transcripts (green) to the left are known to be strongly P-responsive from previous ATH1 studies, whereas the 19 (pink) are not represented on ATH1 arrays and 5 (yellow) are not even annotated. AGIs for these 5 unannotated genes to the right (A, B) are therefore tentative. Expression levels are shown on a double logarithmic scale as 40-ΔC_T_, where ΔC_T_ is the difference between the C_T_ (threshold cycle number) of the respective gene and the reference gene (*UBQ10*; *At4g05320*); therefore 40 equals the expression level of UBQ10. An arbitrary number 40 was selected because RT-qPCR run stops after 40 cycles. (C) Venn diagram showing the number of PRGTs regulated by *PHR1, PHR1PHL1* and *PHO2*.

### Alternative splicing of PRGTs and identification of *miR399* resistant *PHO2*.*2*

Inspection of the genome using splice junction mapper TopHat2 v2.0.9 (Kim et al., 2013) in 100 nt windows and looking for significant changes in coverage within genes or hotspot list identified almost 100 additional PRGTs, many significantly improved annotations and some alternative splicing events (supplemental Table S6 and S7). GO analysis of the 44 alternately spliced genes revealed chromatin-binding or -regulatory protein as the most enriched GO category followed by gene-specific transcriptional regulator, membrane traffic protein, metabolite interconversion enzyme, nucleic acid metabolism protein, protein modifying enzyme, transmembrane signal receptor, and transporter (supplemental Figure S9). Among these alternately spliced PRGTs was *PHO2* (*PHOSPHATE 2*), which has a splice form that encodes only about half the transcript by splicing out of the miR399 binding sites, 1^st^, 2^nd^ and 3^rd^ introns as well as 2^nd^ exon of 5’ UTR and 1^st^ exon of the coding sequence (Figure 7A). As *PHO2* is known to play a critical role in Pi-signaling pathway and homeostasis (Bari et al., 2006; Pant et al., 2008), its alternative splicing was further investigated in detail. Interesting splice variants of *PHO2* (*PHO2*.*1 and PHO2*.*2*) were detected using IGV, therefore six pairs of splice form specific primers were designed to verify its existence by PCR and RT-qPCR. Data from the PCR based assay confirmed the existence of *PHO2*.*1* and *PHO2*.*2*, a larger and smaller splice form of *PHO2*, respectively (Figure 7B). Furthermore, transcript abundance of *PHO2* was monitored with splice form specific RT-qPCR primers (Figure 7C, D). *PHO2*.*2* (shorter splice form of *PHO2*) has relatively higher abundance in P-limited shoot, whereas it is almost undetectable in shoots of FN grown plants (Figure 7C) but in roots that form is equally abundant in both conditions. This shows that *PHO2*.*2* level is not affected by *miR399* abundance, which is very high in P-limited roots. Splice form *PHO2*.*1* is more abundant in P-sufficient root compared to P-limited root, this fits with the *miR399* mediated negative regulation of *PHO2* during P-limitation. But in shoot, *miR399* mediated negative regulation of *PHO2* does not take place (Bari et al., 2006; Pant et al., 2008). The splice form *PHO2*.*2* is equally abundant in P-sufficient and P-limited root, fitting with the model that *PHO2*.*2* is resistant to *miR399*-mediated transcript cleavage, as it does not have a *miR399* binding site. Intriguingly, *PHO2*.*2* is undetectable in P-sufficient shoot but more abundant in P-limited shoot (Figure 7A1, D). This suggests a novel mechanism involving negative regulation of *PHO2*.*2* in P-sufficient shoot.

**Figure 7.**
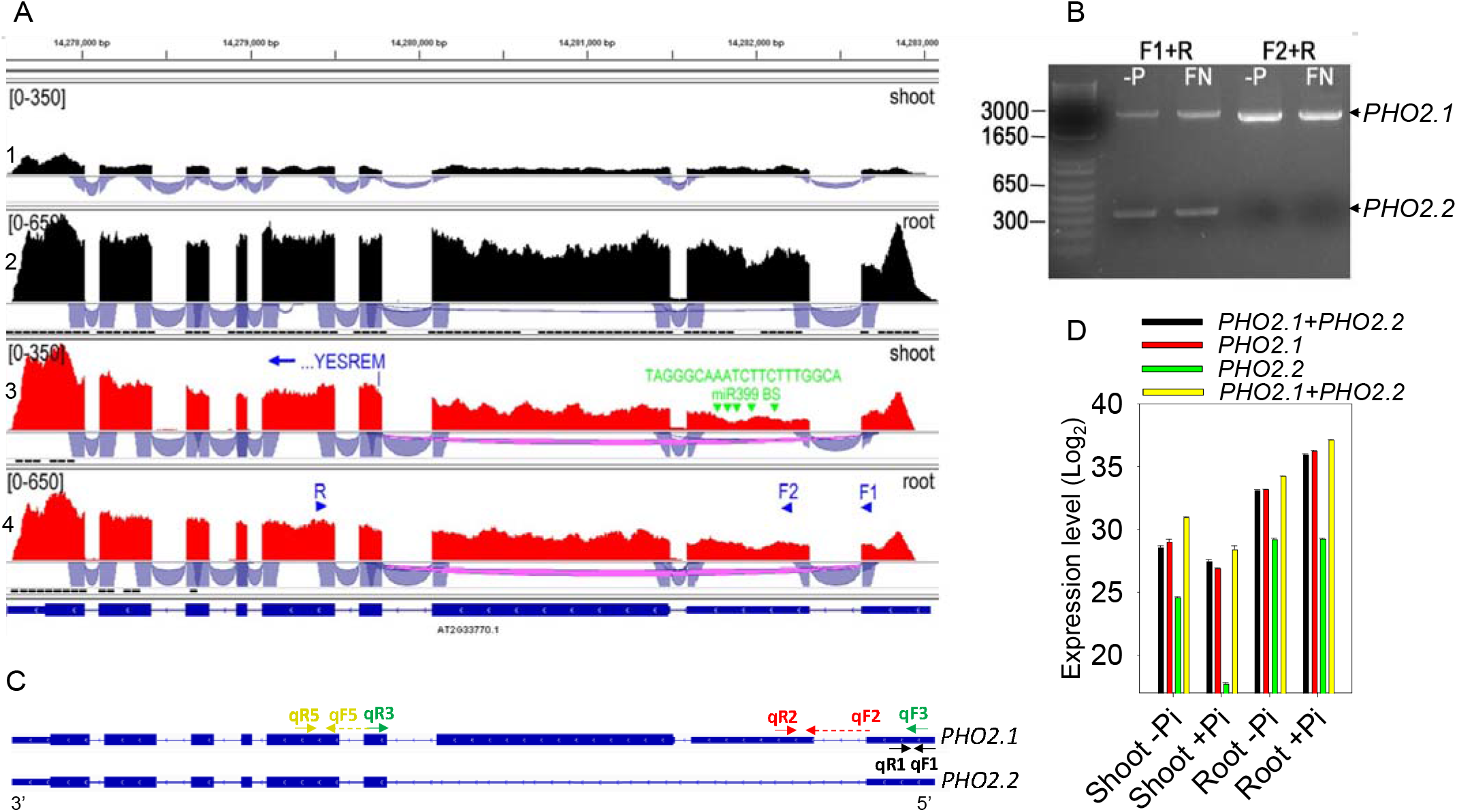
Alternative splicing of *PHO2* during P-limitation. (A) Sashimi plots and splice junction tracks are shown for *PHO2* (*At2g33770*) in P-sufficient (black) and P-limited (red) shoots and roots. The alternative splicing event that leads to elimination of a large portion of *PHO2* including 1^st^, 2^nd^ and 3^rd^ introns, 2^nd^ exon of 5’ UTR and 1^st^ exon of coding sequence is shown in pink in the 3^rd^ and 4^th^ Sashimi plot. *MicroRNA399* binding sites (*miR399* BS; green arrow heads) and their consensus sequence are shown in green in the third plot, as well as the presumable alternative translational start in exon 4 (MERSEY…, blue). (B) Agarose gel detection of *PHO2* splice variants amplified with primers F1+R or F2+R (indicated in blue in the 4^th^ plot) and cDNA produced from roots of P-limited (-P) or P-sufficient (FN, +P) *Arabidopsis* plants. PCR product sizes amplified with F1+R or F2+R are 2460 / 315, or 2286 nucleotides. (C) Schematic representation of *PHO2* splice forms (*PHO2*.*1* and *PHO2*.*2*) showing RT-qPCR primer locations with arrows and dotted arrows for exon-exon junctions. Thick boxes represent exons, thin boxes represent UTRs and lines represent introns.. (D) Bar diagram showing RT-qPCR expression levels for *PHO2*.*1* and *PHO2*.*2* transcripts in shoot and root in +P and -P conditions. The expression levels are shown on a log_2_ scale as 40-ΔCT. The data are average of three replicates ± standard error (SE). Primer pairs qF1+qR1 detects both *PHO2*.*1* and *PHO2*.*2*, qF2+qR2 detects *PHO2*.*1*, qF2+qR3 detects *PHO2*.*2*, and qF4+qR4 detects both *PHO2*.*1* and *PHO2*.*2*.

### Phosphate limitation modulates plant defense via PHR1-PHL1

Phosphate deficiency was previously found to change in the expression of jasmonic acid (JA) biosynthetic (e.g. *AT3G25760*) or signaling genes (e.g. *AT3G12500*) (Morcuende et al., 2007; Bustos et al., 2010) and to induce the jasmonate pathway enhancing resistance to insect herbivory (Khan et al., 2016). JA induction and P starvation also share some common phenotypes, such as growth reduction and anthocyanin accumulation. Therefore, the response of gene transcripts involved in defense responses was analyzed in more detail. *VSP2* expression was ∼10-fold upregulated in leaves, but *JASMONATE-ZIM-DOMAIN PROTEIN 10* (*JAZ10*) and *LIPOXYGENASE 2* (*LOX2*) were not affected. In addition, several genes encoding plant defensins (*At1g75830*/*PDF1*.*1, At2g02120*/*PDF2*.*1, At2g02130*/*PDF2*.*3, AT5G56368, AT2G43535*) and gene transcripts encoding TIR-NBS disease resistance proteins (*AT1G72890, AT3G04210, AT1G17615, AT1G63750*) were strongly induced in leaves and roots during P-limitation. Another defense-related gene transcript with strong P-limitation induction is *AT4G12470* (*AZELAIC ACID INDUCED 1, AZI1*), which is involved in the priming of salicylic acid induction and systemic immunity triggered by pathogens or azelaic acid (Jung et al., 2009). To determine if PHR1-PHL1 is involved in regulation of these plant defense related PRGTs, we analyzed their expression by RT-qPCR. We found that majority of these genes, which encodes DISEASE RESISTANCE PROTEINS, DEFENSIN-LIKE PROTEINS, TN13 and PATHOGENESIS-RELATED 3, were induced by P-limitation and showed attenuated expression in P-limited *phr1* and *phr1-phl1* mutants (Figure 8A).

**Figure 8.**
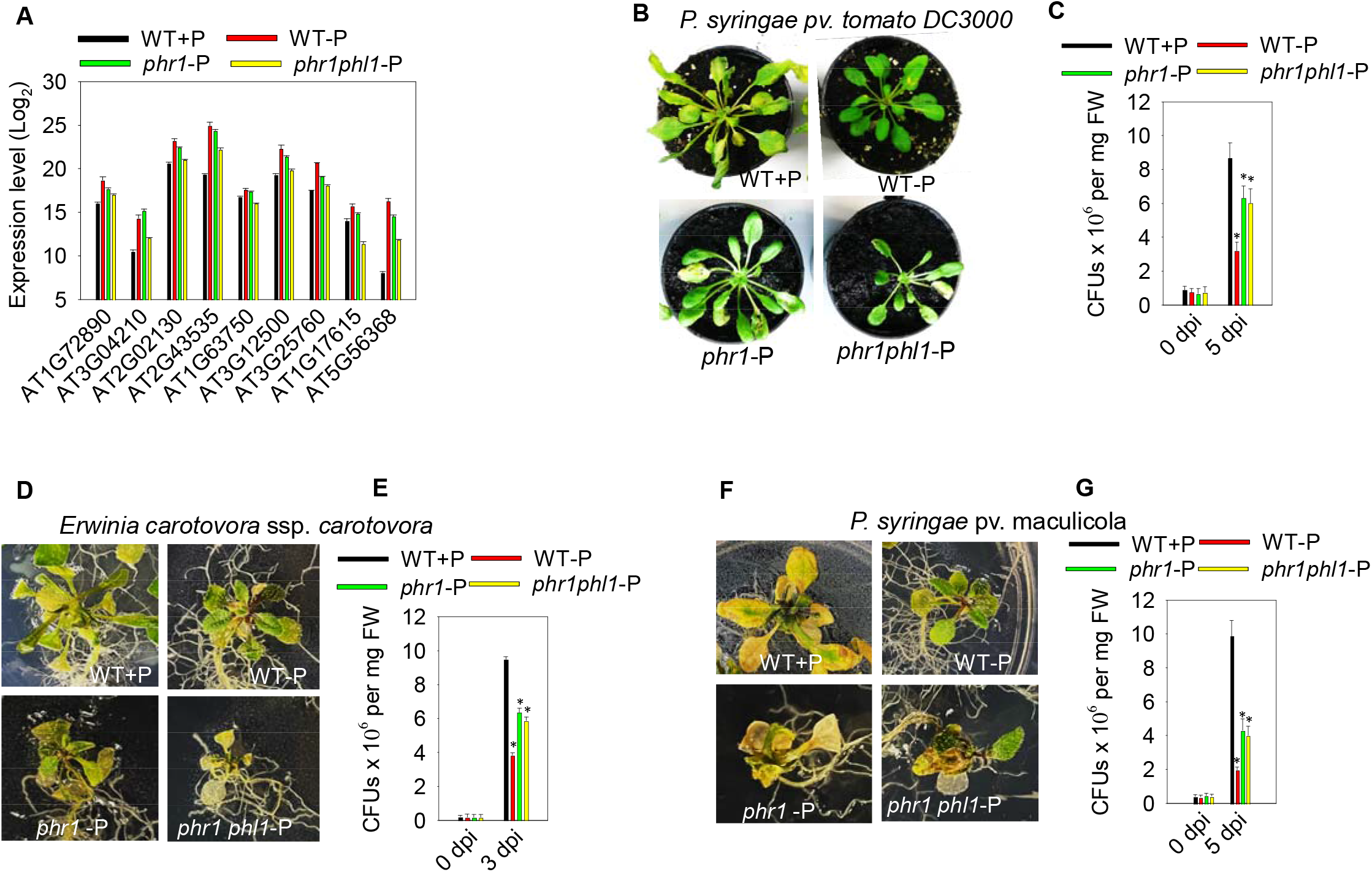
Phosphate limitation modulates plant defense via PHR1-PHL1. (A) RT-qPCR expression analysis of plant defense related genes in wild-type (WT), *phr1* and *phr1phl1* mutants grown under P-limited (-P) condition. Gene expression levels are shown on a Log_2_ scale along the y-axis. Gene expression in WT plants grown under P-sufficient condition (WT+P) was taken as a control. The data are average of three replicates ± standard error (SE). (B) Phenotypic response of 4-week-old *Arabidopsis thaliana* Col-0 grown under short day condition (8:16 light-dark cycle and light intensity 140 µmol m^-2^ s^-1^) at 5 dpi. Plants were spray inoculated with bacterial suspension of *P. syringae* pv. tomato DC3000 at concentration of 1×10^8^ CFU/ml in water containing 0.01% Silwet L-77. Control plants were inoculated with sterile water containing 0.01% Silwet L-77. (C) Quantification of *P. syringae* pv. tomato DC3000. (D) *Arabidopsis* seedlings were flood inoculated with *Erwinia carotovora* ssp. *Carotovora* at concentration of 1.6×10^5^ CFU/ml in water containing 0.01% Silwet L-77 at 3 dpi. (E) Quantification of *Erwinia carotovora* ssp. *carotovora*. (F) *Arabidopsis* seedlings were flood inoculated with *P. syringae* pv. maculicola at concentration of 1.6×10^5^ CFU/ml in water containing 0.01% Silwet L-77 at 5 dpi (G) Quantification of *P. syringae* pv. maculicola. Standard error from six biological replicates is shown as error bars in all graphs and experiments were repeated with similar results. Asterisks is shown for statistically significant difference with Student’s t test (p value ≤ 0.05) when compared to control.

These results suggest that P-limited plants could be more resistant to pathogens. To confirm this hypothesis, we grew *Arabidopsis* (Col-0) plants under P-limitation and P-sufficient conditions. A bacterial spray-based disease assay was performed with a biotrophic pathogen, *Pseudomonas syringae* pv. tomato DC3000 in plants grown in soil. Plants grown in soil or on agar under P-limited conditions show a slightly different developmental phenotype. To see if agar grown plants under P-limited condition are also resistant to pathogens, we used a flood inoculation method (Ishiga et al., 2011; Ishiga et al., 2017) and used a different biotrophic pathogen, *P. syringae* pv. maculicola and a necrotrophic pathogen *Erwinia carotovora* ssp. *carotovora*. The results showed that P-limited plants grown either in soil or on agar were resistant to all the pathogens tested as evidenced by reduced necrotic/chlorotic spots when compared to P-sufficient plants (Figure 8B-G). In addition to the disease symptoms, the bacterial growth curve assay also showed significantly less bacterial accumulation in P-limited *Arabidopsis* plants when compared to P-sufficient plants. Pathogenicity assay also showed that *phr1* -P and *phr1-phl1* -P were more susceptible to these pathogens when compared to wild-type -P plants (Figure 8B-G). These results indicate that P-limitation enhances disease resistance in *Arabidopsis* and is regulated via *PHR1-PHL1*.

### P-limited plants transpire less water compared to P-sufficient plants and is mediated by PHR1-PHL1

We found that *AT4G27920*, which encodes a member of the *PYRABACTIN RESISTANCE* (*PYR*)*/PYR1-like* (*PYL*)*/REGULATORY COMPONENTS OF ABA RECEPTOR* (*RCAR*) family proteins was induced in both shoots and roots during P-limitation. Plant induction of ABA during drought stress is sensed by PYR/PYL/RCAR receptors and these in turn regulate the activity of protein phosphatases 2Cs ABI1/ABI2 and downstream signaling pathway during drought stress (Santiago et al., 2009; Cutler et al., 2010; Yu et al., 2016). *AT1G69260*, which encodes ABI FIVE BINDING PROTEIN (AFP1) was also induced in shoot of P-limited plants.

To further determine the cross talk between P-limitation and drought, we identified the genes confirmed to play role in drought based on literature search in TAIR10 database (www.arabidopsis.org) and found that 11 of these genes were induced by P-limitation by more than 5-fold (Figure 9A). GO term enrichment for these genes revealed that majority of these genes are transcriptional regulators and transporters. Among these, *AT1G21529*, which encodes a DROUGHT INDUCED lncRNA (DRIR), was ∼300 fold induced by P-limitation. Overexpressing *DRIR* in *Arabidopsis* was shown to increase drought tolerance and ABA sensitivity (Qin et al., 2017). *MiR399f*, a highly P-limitation induced gene, also modulates plant responses to ABA and drought (Baek et al., 2016), in addition to its role in phosphate signaling. *AT5G09570* (*AT12CYS-2*), which is highly induced by P-limitation (Figure 9B), encodes a twin CX9C domain protein and loss of both *AT12CYS-1* and *AT12CYS-2* was shown to enhance drought tolerance (Wang et al., 2016). We performed RT-qPCR expression analysis of eight genes common between drought and P-limitation and found that seven out of eight genes were regulated via PHR1-PHL1 transcription factors (Figure 9B). However, *AT12CYS-2*, which is a negative regulator of drought tolerance, was also induced in *phr1* or *phr1-phl1* mutants during P-limitation.

**Figure 9.**
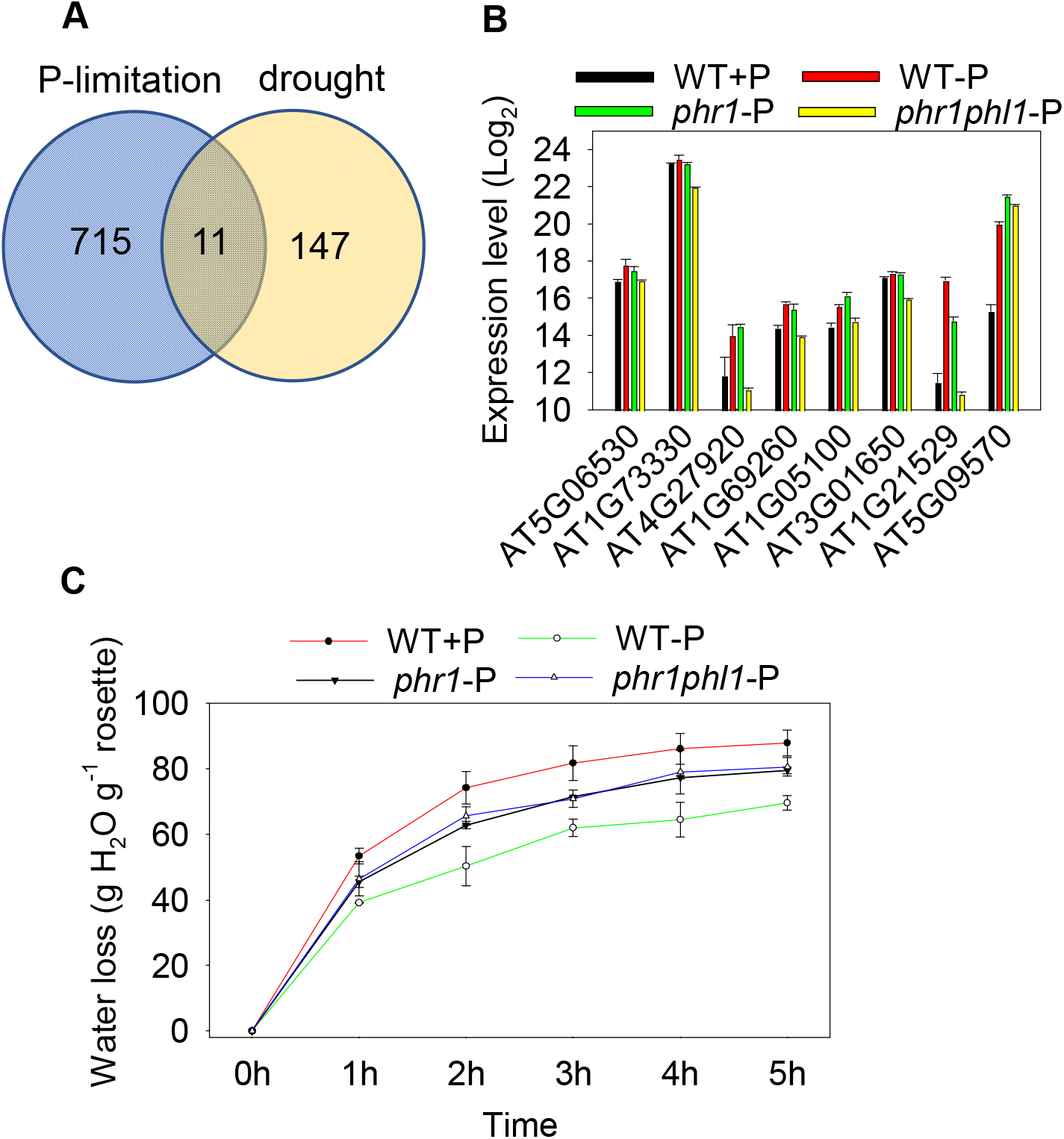
P-limited plants show enhanced drought tolerance that is mediated by PHR1-PHL1. (A) Venn diagram of genes known to play role in drought based on literature search (www.arabidopsis.org) and annotated genes induced by P-limitation (≥ 5-fold in shoot and/or root). (B) RT-qPCR expression profiling of eight drought related genes in WT, *phr1* and *phr1phl1* seedlings grown under P-limited condition. Gene expression levels are shown on a Log_2_ scale along the y-axis. *Arabidopsis* seedlings grown under P-sufficient condition (WT+P) was taken as control. The data are average of three replicates ± standard error (SE). (C) Water loss assay performed in WT, *phr1* and *phr1phl1* rosettes grown under P-sufficient (+P) or P-limitation (- P) condition. The data are average of eight biological replicates ± SE representing two independent experiments.

Based on all these results, we hypothesized that P-limitation can enhance drought tolerance in plants. Water loss assay in the detached leaves of *Arabidopsis* WT, *phr1* and *phr1-phl1* mutants was performed and it confirmed that P-limited plants lost the least amount of water compared to P-sufficient plants, while P-limited *phr1* and *phr1-phl1* mutants lost more than WT-P (Figure 9C). This suggests that P-limitation may enhance plant drought tolerance and is mediated via PHR1-PHL1.

## DISCUSSION

Owing to the unbiased nature and relatively high sensitivity of the sequencing approach, our RNA-Seq analysis revealed a larger number of novel, previously undescribed P-limitation responsive gene transcripts (PRGTs). These include novel annotated and unannotated genes and alternative splice forms. The majority of P-limitation induced genes showed reversion after 30 minutes and 3 hours of P addition to P-limited plants, and were regulated by *PHR1, PHL1* and *PHO2*. Our results also showed the involvement of many novel PRGTs in the cross-talk with other stresses including enhanced plant defense and drought tolerance. We have also shown that PHR1-PHL1 mediates the P-limitation induced plant defense and drought tolerance.

We identified more than 1,000 PRGTs that are ≥ 5-fold induced or repressed in shoot and/or root during P-limitation. Of these, ∼75% PRGTs were up-regulated and ∼25% were repressed, which is similar to 3:1 ratio of induced vs. repressed PRGTs in the ATH1 study of Bustos et al., (2010). Importantly, ∼60% of the PRGTs detected in this study were not previously reported in ATH1 experiments, although 26% have an associated ATH1 gene identifier. Some of the differences in the number of PRGTs in different studies could be due to biological variation between experiments (plant, age etc.) or technical differences. For example, difference in P-response to Bustos et al. 2010 experiment could be due to lack of specificity and cross-hybridization of ATH1 probe sets (Figure S3 and S4) (e.g., 259221_s_at for *AT3G03540*) with RNA from a highly homologous locus (i.e., *AT3G03530*). The probe set (259221_s_at) perfectly matches *AT3G03530*, and hence picks up the signal of *AT3G03530*, and attributes it to *AT3G03540*. RNA-Seq however indicates that only *AT3G03530* is expressed (Figure S10), as the two homologous gene transcripts are still sufficiently different (86% identity) for unambiguous read mapping. Experimental variations are also commonly observed among studies (Lee et al., 2020). There might also be genes that were simply not detected by StringTie for example *miR399a* (*AT1G29265*), *At2g34210, AT4G38080* or *AT2G41070* because their transcripts were fused with those from neighboring genes (supplemental Figure S11). Also, there are a few examples where the Sashimi plot contradicts the StringTie calculation and confirms the P-responsiveness of some genes identified by Bustos et al., 2010 (e.g. *AT4G38320*). Although, Cufflinks (Trapnell et al., 2010) and StringTie (Pertea et al., 2015) are currently the most widely used genome-guided transcriptome assemblers and StringTie correctly assembles 20% more transcripts than Cufflinks (Pertea et al., 2015). We found that none of these transcript assemblers are perfect, and both miss some transcripts; however, StringTie was better at assembling more transcripts and therefore led to the identification of more PRGTs in this study. *AT1G76430/Pht1*.*9* was not identified by StringTie and is an example showing that also HiSat StringTie doesn’t work perfectly. Therefore, one should not completely rely on assembler output and visual inspection in IGV is important for robustness.

Our data showed a link between P-limitation and ion homeostasis. *AT2G43500/NLP8*, a member of the RWP-RK regulator family containing the octicosapeptide/Phox/Bem1p domain, was found to be induced during P-limitation in shoots. Its homolog NLP7 modulates nitrate sensing and metabolism, and *nlp7* mutants show features of nitrogen-starved plants and are tolerant to drought stress (Castaings et al., 2009). The RWP-RK domain of NLPs was shown to bind to nitrate-responsive elements (Konishi and Yanagisawa, 2013). Thus, NLPs might provide a novel link between P- and N-signaling in plants. *AT1G13300/NIGT1*.*4/HRS1*, which integrate nitrate and phosphate signals at the *Arabidopsis* root tip (Medici et al., 2015), was more P-limitation induced in our study than reported previously in *Arabidopsis* ATH1 gene chip studies. Many PRGTs encoding high-affinity nitrate transporters were strongly repressed in roots, suggesting that P-limitation decreases plants capacity for nitrate uptake. Previous ATH1 gene chip studies identified several PRGTs encoding phosphate transporters, and some potassium, sulfur and nitrate transporters (Misson et al., 2005; Morcuende et al., 2007; Bustos et al., 2010). However, P-limitation induction of *PHO1*.*H10* in shoot was not reported previously. Our RNA-seq study revealed several novel PRGTs encoding other cation/anion transporters, peptide transporters, vacuolar iron transporters, sugar, ABC, MtN21 and MATE transporters. Interestingly, five Nodulin MtN21-like/UMAMIT (USUALLY MULTIPLE ACIDS MOVE IN AND OUT TRANSPORTERS) transporter family members were induced during P-limitation. As the concentration of amino acids are increased during P-limitation (Morcuende et al., 2007; Pant et al., 2015), induction of these *UMAMIT* genes, mainly in P-limited shoots, suggest their involvement in importing amino acids from xylem which could contribute to higher amino acid accumulation in the leaves during P-limitation.

We identified the genes linking P-limitation to hormone signaling. *AT1G68320*/*MYB62* was found among the P-responsive TFGTs. *MYB62* is involved in regulation of phosphate starvation responses and gibberellic acid biosynthesis (Devaiah et al., 2009). Additional links between P-limitation and hormone signaling are provided by P-responsive TFGTs such as *ETHYLENE-INSENSITIVE3-LIKE 2* (*EIL2*), ethylene response factors (*ERF22, RAP2*.*6, ERF72* and *SHN3*), *ABA REPRESSOR1* (*ABR1*), and *IAA5*. Many P-limitation induced transcriptional regulators such as *PRR9, CCA1, LHY* and *LCL5* are involved in clock function (Schaffer et al., 1998; Wang and Tobin, 1998; Matsushika et al., 2000; Farinas and Mas, 2011). CCA1 functions synergistically with LHY in regulating circadian rhythms of *Arabidopsis*, while PRR9 is an essential component of the temperature-sensitive circadian system and acts as transcriptional repressor of CCA1 and LHY (Nakamichi et al., 2010). LCL5 is involved in the regulation of circadian clock by modulating the pattern of histone 3 (H3) acetylation (Farinas and Mas, 2011). These findings suggest that P-limitation affects circadian rhythms via induction of these central clock TFs.

P-limited plants develop more and longer root hairs when compared to P-sufficient plants (Raghothama, 1999). R3-type MYB genes such as *ENHANCER OF TRY AND CPC1* (*AT1G01380/ETC1*) and *CAPRICE-LIKE* (*AT4G01060/CPL3/ETC3*) are known as key regulators of plant root-hair development (Kirik et al., 2004; Tominaga-Wada and Wada, 2014; Wada et al., 2015). We found that these *CPC* family genes were induced in the roots of P-limited plants suggesting their important role in the regulation of root hair development during P-limitation. In addition, the present study identified many novel P-limitation induced, long-noncoding RNAs, microRNAs, transcripts that encode proteins involved in signaling such as small peptides, receptor kinases and MAP kinases, and components of protein degradation machinery and proteins involved in post-translational protein modification including kinases and phosphatases. Several nutrient responsive peptides were identified in *Medicago truncatula* (de Bang et al., 2017) and peptides are known to play key role in plant growth and stress response (Murphy et al., 2012; Smith et al., 2020; Hu et al., 2021) signifying the importance of these novel peptides.

We identified more than 100 P-limitation inducible unannotated transcripts which were not reported previously and the majority of those were lncNAs. LncRNAs that regulate gene expression by chromatin remodeling or transcriptional interference, by acting as microRNA target mimics s, antisense RNAs, alter the stability and translation of mRNAs (Franco-Zorrilla et al., 2004; Statello et al., 2021). Some of these transcripts can also encode small functional peptides (Choi et al., 2019) and it will be interesting to investigate the function and small peptide coding potential of these lncRNAs. We identified and confirmed that one of these lncRNAs was a microRNA that targets a protein kinase involved in signal transduction (Shiu and Bleecker, 2001) and an F-box protein involved in brassinosteroid mediated signaling pathway (Zhu et al., 2017), suggesting an important role of this novel microRNA in plant signaling.

We identified several alternatively spliced transcripts during P-limited conditions. Here, we report for the first time about the alternative splicing of *PHO2* to form a shorter splice form (*PHO2*.*2*) that does not contain *miR399* binding sites, therefore escapes *miR399* mediated target degradation. *PHO2* contains five *miR399* binding sites in its 5’ UTR and is negatively regulated by *miR399* via transcript degradation (Bari et al., 2006). During P-limitation, *miR399* is induced in the shoot and moves to the root where it negatively regulates *PHO2* by transcript degradation (Pant et al., 2008). Our finding of downregulation of *PHO2*.*1* in root -P compared to root +P agrees with previous findings of *miR399* mediated transcript degradation of *PHO2* in the root (Pant et al., 2008). However, *PHO2*.*2* is not altered in root -P vs root +P but it is strongly decreased in shoot +P compared to shoot -P. This suggests distinct regulation of *PHO2*.*2* in shoot and root. *PHO2*.*1* encodes a ubiquitin-conjugating E2 enzyme (UBC24) with a large N-terminal regions with no known domains (Bari et al. 2006 and present study). We found that a large part of this N-terminal region is spliced out in *PHO2*.*2*. Interestingly, the catalytic UBC domain of PHO2, which is located towards C-terminal is intact in PHO2.2. PHO2 has high homology to human E2/E3 hybrid enzyme (UBE2O), which is known to play diverse roles in regulating protein ubiquitination, cellular functions, and disease onset (Kraft et al., 2005; Ullah et al., 2019; Liu et al., 2020). Although, functional characterization of different *PHO2* splice forms is beyond the scope of this paper, future studies might reveal a novel mechanism of *PHO2*.*2* regulation to maintain P signaling and homeostasis in plants.

We found that the genes related to fatty acid (FAs) metabolism such as *AT4G01950* (*glycerol-3-phosphate acyltransferase*), *AT4G12470* (*AZI1*), *AT2G01180* (*phosphatidate phosphatase galactolipids*) were induced during P-limitation. It is known from previous studies that *AZI1* and FAs are involved in plant systemic immunity to pathogens (Jung et al., 2009; Kachroo and Kachroo, 2009). Many genes for plant defensins and TIR-NBS (Toll/interleukin receptor nucleotide-binding site Leucine-rich repeat) disease resistance proteins involved in plant disease resistance (Thomma et al., 2003; McHale et al., 2006; Nandety et al., 2013) were found to be induced by P-limitation. These factors could contribute to the pathogen resistance in P-limited plants. We were able to experimentally confirm that P-limited plants are more resistant to pathogens and this response is mediated by PHR1-PHL1. TRXL1, which is involved in plant thermotolerance and disease resistance (Pant et al., 2020) is also induced by P-limitation providing another link between P-limitation and other stresses. Previously, we showed that P-limitation leads to increases in plant secondary metabolites including glucosinolates (Pant et al., 2015). Glucosinolates are antimicrobial compound and increase plant resistance to pathogens (Frerigmann et al., 2012). P-limitation induced increase in plant resistance against pathogens can also be attributed to increased glucosinolates and other secondary metabolites in addition to the increased expression of defense related genes.

We found that *PYR/PYL/RCAR* was induced by P-limitation suggesting it as a common component for cross-talk between drought and P-limitation. PYR/PYL/RCAR family proteins mediate ABA-dependent regulation of protein phosphatase 2Cs (Fujii et al., 2009; Ma et al., 2009; Park et al., 2009) and downstream signaling pathway during drought stress (Vlad et al., 2009). We found that many drought related genes were induced by P-limitation and were regulated by *PHR1-PHL1*. PHR1 (PHOSPHATE RESPONSE 1) is a master regulator of P-starvation response in plants (Rubio et al., 2001; Bari et al., 2006; Bustos et al., 2010; Pant et al., 2015; Pant et al., 2015). We found that several drought and defense related genes were regulated by *PHR1-PHL1* suggesting PHR1-PHL1 is involved in cross-talk between drought/defense and plant Pi homeostasis. Our experimental verification of P-limited plants being drought tolerant or pathogen resistance can be useful to develop new management strategies to protect crop from multiple stress by creating mild P-limited condition. In addition to PHR1-PHL1, we identified that a *DROUGHT INDUCED LONG NON-CODING RNA* (*DRIR*), which is known to enhances plant drought tolerance (Qin et al., 2017) and could potentially be involved in cross-talk between P-limitation and drought stress signaling.

Identification of unannotated and annotated novel P-responsive genes involved in regulatory and signaling function, noncoding RNAs, small peptides, novel P-responsive splice forms including *miR399* resistant *PHO2*.*2*, transporters, genes involved in root hair formation and defense add to our understanding of P-signaling (Figure 10). Furthermore, the involvement of PRGTs in hormone signaling, drought and defense pathway entails cross-talk between P-limitation with other stress pathways including plant defense. Regulation of the majority of PRGTs reversibly by P-availability and their regulation by *PHR1, PHL1* and *PHO2* add new potential players to the P-signaling network. Novel insights provided by this study can be useful for translational research to develop genetically enhanced stress resilient crop varieties that can grow in P-limitation conditions. In addition, these results could inform more efficient management strategies.

**Figure 10.**
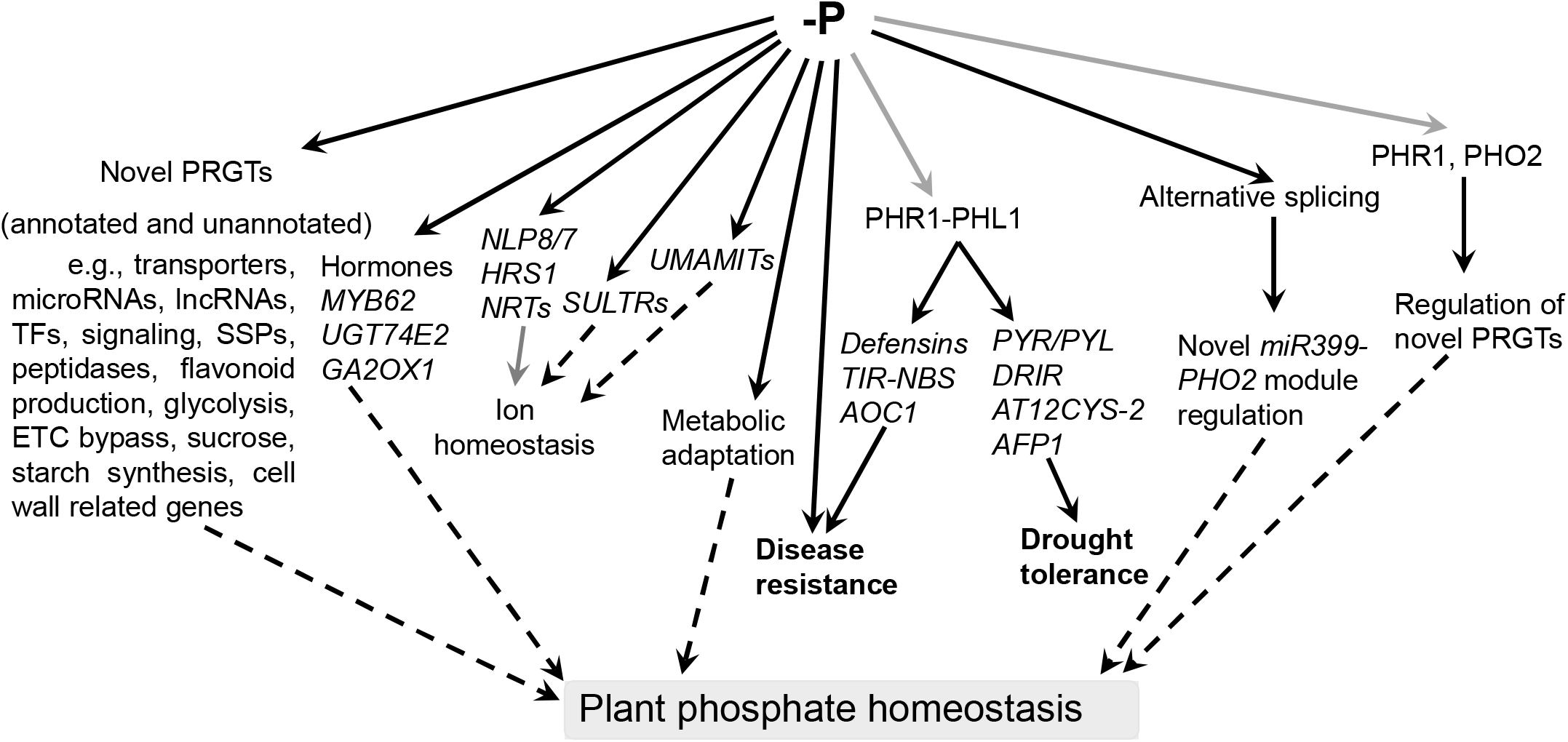
P-limitation responsive transcriptomics identifies novel and unannotated P-responsive genes, alternative splice forms and cross-talk to abiotic and biotic stress. Phosphate limitation (-P) regulates expression of several novel PRGTs, hormone related genes (*MYB62, UGT74E2, GA2OX1*), *UMAMITs, NLP8/7, HRS1, NRTs, SULTRs*, disease resistance genes (*TIR-NBS, AOC1, LCR67/70*), drought related genes (*PYR/PYL*, ABA, PHR1), alternative splicing of PRGTs including *PHO2*, regulation of novel PRGTs by PHR1 and PHO2. Major category of the novel annotated PRGTs includes transporters, microRNAs, long non-coding RNAs (lncRNAs), transcription factors (TFs), genes involved in signaling and small secreted peptide (SSP), peptidases, flavonoid production, glycolysis, mitochondrial electron transport chain (ETC) bypass, hormone metabolism, sucrose and starch synthesis, cell wall metabolism. Gray colors for arrows and letters represent the known information from the previous literatures and black arrows and letters show the novel information identified in the present study. *UGT74E2, URIDINE DIPHOSPHATE GLYCOSYLTRANSFERASE 74E2*; *GA2OX1, GIBBERELLIN 2-OXIDASE*; *NLP, NIN LIKE PROTEIN*; *HRS1, HYPERSENSITIVITY TO LOW PI-ELICITED PRIMARY ROOT SHORTENING 1*; *NRT, NITRATE TRANSPORTERS*; *SULTR, SULPHATE TRANSPORTER*; *UMAMIT, USUALLY MULTIPLE ACIDS MOVE IN AND OUT TRANSPORTERS*; *TIR-NBS, TRUNCATED TOLL/INTERLEUKIN-1 RECEPTOR– NUCLEOTIDE-BINDING SEQUENCE*; *AOC1, ALLENE OXIDE CYCLASE 1*; *PYR/PYL, PYRABACTIN RESISTANCE*)*/PYL*(*PYR1-like*); *DRIR, DROUGHT INDUCED LONG NCRNA* (*DRIR*); *AT12CYS-2, TWIN CYSTEINE PROTEINS*; *AFP1, ABI FIVE BINDING PROTEIN*; *PHR1, PHOSPHATE STARVATION RESPONSE 1*; *PHO2, PHOSPHATE 2*.

## Supplementary data

Supplemental Figure S1. Aspect and quality control of Arabidopsis materials

Supplemental Figure S2. Venn diagram showing the phosphate responsive gene transcripts (PRGTs)

Supplemental Figure S3. ATH1 probe sets do not properly detect some of the actual gene transcripts

Supplemental Figure S4. Re-annotation of AT1G30550 based on RNA-seq data

Supplemental Figure S5. Confirmation of P-status responsive gene transcripts by RT-qPCR

Supplemental Figure S6. Sashimi plots of unannotated highly P-status responsive transcripts.

Supplemental Figure S7. Additional information on annotation and sequence information for the novel PRGTs that are currently unannotated in TAIR10 database.

Supplemental Figure S8. Inter-study comparison of PRGTs in shoots.

Supplemental Figure S9. GO analysis of 44 alternately spliced genes.

Supplemental Figure S10. Sashimi plots for selected strongly P-status responsive gene transcripts.

Supplemental Figure S11. Sashimi plot for miR399a (AT1G29265) as an example of PRGTs missed by RNA-seq analysis pipeline.

Supplemental Table S1. RNA-seq analysis showing gene abundance, top 50 genes, phosphate regulated gene transcripts (PRGTs), unannotated P-responsive transcripts and transcription factors.

Supplemental Table S2. Transcription factor genes with P-status responsive transcript abundance as detected in this study using RNA-seq and comparison with previous studies

Supplemental Table S3 Comparison of 11 lncRNAs (no ATH1 ID) in Yuan et al. 2016 and present study

Supplemental Table S4 Comparison of P-response of unannotated transcriptional units with Yuan et al., 2016

Supplemental Table S5: PHR1 binding site (P1BS) in promoters in novel PRGTs

Supplemental Table S6. Alternative splicing of PRGTs

Supplemental Table S7. Examples of improved annotation based on visualization of RNA-seq data in IGV (integrated genome viewer)

Supplemental Table S8. DNA oligos used in the study

## Author contributions

WRS: conceptualization for RNA-seq; PP, BP, KM: conceptualization for remaining part; PP and BP: methodology; NK: processed RNA-seq raw data; WRS, PP and BP: formal analysis; WRS, PP and BP: investigation; WRS, RA and KM: resources; WRS, BP and PP: data curation; WRS, PP and BP: writing - original draft; WRS, PP, BP, RA and KM: writing - review & editing; WRS, PP and BP: visualization; WRS and KM: funding acquisition.

## References

Baek D, Chun HJ, Kang S, Shin G, Park SJ, Hong H, Kim C, Kim DH, Lee SY, Kim MC, Yun DJ (2016) A Role for Arabidopsis miR399f in Salt, Drought, and ABA Signaling. Mol Cells 39: 111–118

Baek D, Chun HJ, Yun DJ, Kim MC (2017) Cross-talk between Phosphate Starvation and Other Environmental Stress Signaling Pathways in Plants. Mol Cells 40: 697–705

Bandyopadhyay S, Mitra R (2009) TargetMiner: microRNA target prediction with systematic identification of tissue-specific negative examples. Bioinformatics 25: 2625–2631

Bari R, Pant BD, Stitt M, Scheible WR (2006) PHO2, microRNA399, and PHR1 define a phosphate-signaling pathway in plants. Plant Physiology 141: 988–999

Bolger AM, Lohse M, Usadel B (2014) Trimmomatic: a flexible trimmer for Illumina sequence data. Bioinformatics 30: 2114–2120

Bustos R, Castrillo G, Linhares F, Puga MI, Rubio V, Perez-Perez J, Solano R, Leyva A, Paz-Ares J (2010) A central regulatory system largely controls transcriptional activation and repression responses to phosphate starvation in Arabidopsis. PLoS genetics 6: e1001102

Castaings L, Camargo A, Pocholle D, Gaudon V, Texier Y, Boutet-Mercey S, Taconnat L, Renou JP, Daniel-Vedele F, Fernandez E, Meyer C, Krapp A (2009) The nodule inception-like protein 7 modulates nitrate sensing and metabolism in Arabidopsis. Plant J 57: 426–435

Choi SW, Kim HW, Nam JW (2019) The small peptide world in long noncoding RNAs. Brief Bioinform 20: 1853–1864

Cordell D, White S (2014) Life’s Bottleneck: Sustaining the World’s Phosphorus for a Food Secure Future. Annual Review of Environment and Resources 39: 161–188

Cutler SR, Rodriguez PL, Finkelstein RR, Abrams SR (2010) Abscisic acid: emergence of a core signaling network. Annu Rev Plant Biol 61: 651–679

Darcy JL, Schmidt SK, Knelman JE, Cleveland CC, Castle SC, Nemergut DR (2018) Phosphorus, not nitrogen, limits plants and microbial primary producers following glacial retreat. Sci Adv 4: eaaq0942

de Bang TC, Lundquist PK, Dai X, Boschiero C, Zhuang Z, Pant P, Torres-Jerez I, Roy S, Nogales J, Veerappan V, Dickstein R, Udvardi MK, Zhao PX, Scheible W-R (2017) Genome-Wide Identification of Medicago Peptides Involved in Macronutrient Responses and Nodulation Plant Physiology 175: 1669–1689

Devaiah BN, Madhuvanthi R, Karthikeyan AS, Raghothama KG (2009) Phosphate starvation responses and gibberellic acid biosynthesis are regulated by the MYB62 transcription factor in Arabidopsis. Molecular Plant 2: 43–58

Farinas B, Mas P (2011) Functional implication of the MYB transcription factor RVE8/LCL5 in the circadian control of histone acetylation. Plant J 66: 318–329

Franco-Zorrilla JM, Gonzalez E, Bustos R, Linhares F, Leyva A, Paz-Ares J (2004) The transcriptional control of plant responses to phosphate limitation. Journal of Experimental Botany 55: 285–293

Franco-Zorrilla JM, Valli A, Todesco M, Mateos I, Puga MI, Rubio-Somoza I, Leyva A, Weigel D, Garcia JA, Paz-Ares J (2007) Target mimicry provides a new mechanism for regulation of microRNA activity. Nat Genet 39: 1033–1037

Fredeen AL, Rao IM, Terry N (1989) Influence of Phosphorus Nutrition on Growth and Carbon Partitioning in Glycine max. Plant Physiol 89: 225–230

Frerigmann H, Bottcher C, Baatout D, Gigolashvili T (2012) Glucosinolates are produced in trichomes of Arabidopsis thaliana. Front Plant Sci 3: 242

Fujii H, Chinnusamy V, Rodrigues A, Rubio S, Antoni R, Park SY, Cutler SR, Sheen J, Rodriguez PL, Zhu JK (2009) In vitro reconstitution of an abscisic acid signalling pathway. Nature 462: 660–664

Holford ICR (1997) Soil phosphorus: its measurement, and its uptake by plants. Soil Research 35: 227–240

Hou E, Luo Y, Kuang Y, Chen C, Lu X, Jiang L, Luo X, Wen D (2020) Global meta-analysis shows pervasive phosphorus limitation of aboveground plant production in natural terrestrial ecosystems. Nat Commun 11: 637

Hu XL, Lu H, Hassan MM, Zhang J, Yuan G, Abraham PE, Shrestha HK, Villalobos Solis MI, Chen JG, Tschaplinski TJ, Doktycz MJ, Tuskan GA, Cheng ZM, Yang X (2021) Advances and perspectives in discovery and functional analysis of small secreted proteins in plants. Hortic Res 8: 130

Ishiga Y, Ishiga T, Ichinose Y, Mysore KS (2017) Pseudomonas syringae Flood-inoculation Method in Arabidopsis. Bio Protoc 7: e2106

Ishiga Y, Ishiga T, Uppalapati SR, Mysore KS (2011) Arabidopsis seedling flood-inoculation technique: a rapid and reliable assay for studying plant-bacterial interactions. Plant Methods 7: 32

Itaya K, Ui M (1966) A new micromethod for the colorimetric determinationof inorganicphosphate. Clinica Chimica Acta 14: 361–366

Jung HW, Tschaplinski TJ, Wang L, Glazebrook J, Greenberg JT (2009) Priming in systemic plant immunity. Science 324: 89–91

Kachroo A, Kachroo P (2009) Fatty Acid-derived signals in plant defense. Annu Rev Phytopathol 47: 153–176

Khan GA, Vogiatzaki E, Glauser G, Poirier Y (2016) Phosphate Deficiency Induces the Jasmonate Pathway and Enhances Resistance to Insect Herbivory. Plant Physiol 171: 632–644

Kim D, Langmead B, Salzberg SL (2015) HISAT: a fast spliced aligner with low memory requirements. Nat Methods 12: 357–360

Kim D, Pertea G, Trapnell C, Pimentel H, Kelley R, Salzberg SL (2013) TopHat2: accurate alignment of transcriptomes in the presence of insertions, deletions and gene fusions. Genome Biology 14

Kirik V, Simon M, Huelskamp M, Schiefelbein J (2004) The ENHANCER OF TRY AND CPC1 gene acts redundantly with TRIPTYCHON and CAPRICE in trichome and root hair cell patterning in Arabidopsis. Dev Biol 268: 506–513

Konishi M, Yanagisawa S (2013) Arabidopsis NIN-like transcription factors have a central role in nitrate signalling. Nature Communications 4: 1617

Kraft E, Stone SL, Ma L, Su N, Gao Y, Lau OS, Deng XW, Callis J (2005) Genome analysis and functional characterization of the E2 and RING-type E3 ligase ubiquitination enzymes of Arabidopsis. Plant Physiol 139: 1597–1611

Lee AJ, Park Y, Doing G, Hogan DA, Greene CS (2020) Correcting for experiment-specific variability in expression compendia can remove underlying signals. Gigascience 9

Li H, Handsaker B, Wysoker A, Fennell T, Ruan J, Homer N, Marth G, Abecasis G, Durbin R, Genome Project Data Processing S (2009) The Sequence Alignment/Map format and SAMtools. Bioinformatics 25: 2078–2079

Li L, Liu C, Lian X (2010) Gene expression profiles in rice roots under low phosphorus stress. Plant Mol Biol 72: 423–432

Liu X, Ma F, Liu C, Zhu K, Li W, Xu Y, Li G, Niu Z, Liu J, Chen D, Li Z, Fu Y, Qian C (2020) UBE2O promotes the proliferation, EMT and stemness properties of breast cancer cells through the UBE2O/AMPKalpha2/mTORC1-MYC positive feedback loop. Cell Death Dis 11: 10

Lynch J (1995) Root Architecture and Plant Productivity. Plant Physiol 109: 7–13

Ma Y, Szostkiewicz I, Korte A, Moes D, Yang Y, Christmann A, Grill E (2009) Regulators of PP2C phosphatase activity function as abscisic acid sensors. Science 324: 1064–1068

Matsushika A, Makino S, Kojima M, Mizuno T (2000) Circadian waves of expression of the APRR1/TOC1 family of pseudo-response regulators in Arabidopsis thaliana: insight into the plant circadian clock. Plant Cell Physiol 41: 1002–1012

McHale L, Tan X, Koehl P, Michelmore RW (2006) Plant NBS-LRR proteins: adaptable guards. Genome Biol 7: 212

Medici A, Marshall-Colon A, Ronzier E, Szponarski W, Wang RC, Gojon A, Crawford NM, Ruffel S, Coruzzi GM, Krouk G (2015) AtNIGT1/HRS1 integrates nitrate and phosphate signals at the Arabidopsis root tip. Nature Communications 6

Misson J, Raghothama KG, Jain A, Jouhet J, Block MA, Bligny R, Ortet P, Creff A, Somerville S, Rolland N, Doumas P, Nacry P, Herrerra-Estrella L, Nussaume L, Thibaud MC (2005) A genome-wide transcriptional analysis using Arabidopsis thaliana Affymetrix gene chips determined plant responses to phosphate deprivation. Proceedings of the National Academy of Sciences of the United States of America 102: 11934–11939

Morcuende R, Bari R, Gibon Y, Zheng W, Pant BD, Blasing O, Usadel B, Czechowski T, Udvardi MK, Stitt M, Scheible WR (2007) Genome-wide reprogramming of metabolism and regulatory networks of Arabidopsis in response to phosphorus. Plant, Cell & Environment 30: 85–112

Murphy E, Smith S, De Smet I (2012) Small signaling peptides in Arabidopsis development: how cells communicate over a short distance. Plant Cell 24: 3198–3217

Nakamichi N, Kiba T, Henriques R, Mizuno T, Chua NH, Sakakibara H (2010) PSEUDO-RESPONSE REGULATORS 9, 7, and 5 are transcriptional repressors in the Arabidopsis circadian clock. Plant Cell 22: 594–605

Nandety RS, Caplan JL, Cavanaugh K, Perroud B, Wroblewski T, Michelmore RW, Meyers BC (2013) The role of TIR-NBS and TIR-X proteins in plant basal defense responses. Plant Physiol 162: 1459–1472

Ohkubo Y, Tanaka M, Tabata R, Ogawa-Ohnishi M, Matsubayashi Y (2017) Shoot-to-root mobile polypeptides involved in systemic regulation of nitrogen acquisition. Nat Plants 3: 17029

Pant BD, Buhtz A, Kehr J, Scheible WR (2008) MicroRNA399 is a long-distance signal for the regulation of plant phosphate homeostasis. The Plant Journal 53: 731–738

Pant BD, Burgos A, Pant P, Cuadros-Inostroza A, Willmitzer L, Scheible WR (2015) The transcription factor PHR1 regulates lipid remodeling and triacylglycerol accumulation in Arabidopsis thaliana during phosphorus starvation. J Exp Bot 66: 1907–1918

Pant BD, Burgos A, Pant P, Cuadros A, Willmitzer L, Scheible W (2015) The transcription factor PHR1 regulates lipid remodeling and triacylglycerol accumulation in Arabidopsis thaliana during phosphorus starvation. Journal of Experimental Botany

Pant BD, Musialak-Lange M, Nuc P, May P, Buhtz A, Kehr J, Walther D, Scheible WR (2009) Identification of nutrient-responsive Arabidopsis and rapeseed microRNAs by comprehensive real-time polymerase chain reaction profiling and small RNA sequencing. Plant Physiology 150: 1541–1555

Pant BD, Oh S, Lee HK, Nandety RS, Mysore KS (2020) Antagonistic Regulation by CPN60A and CLPC1 of TRXL1 that Regulates MDH Activity Leading to Plant Disease Resistance and Thermotolerance. Cell Rep 33: 108512

Pant BD, Pant P, Erban A, Huhman D, Kopka J, Scheible WR (2015) Identification of primary and secondary metabolites with phosphorus status-dependent abundance in Arabidopsis, and of the transcription factor PHR1 as a major regulator of metabolic changes during phosphorus limitation. Plant Cell and Environment 38: 172–187

Park SY, Fung P, Nishimura N, Jensen DR, Fujii H, Zhao Y, Lumba S, Santiago J, Rodrigues A, Chow TF, Alfred SE, Bonetta D, Finkelstein R, Provart NJ, Desveaux D, Rodriguez PL, McCourt P, Zhu JK, Schroeder JI, Volkman BF, Cutler SR (2009) Abscisic acid inhibits type 2C protein phosphatases via the PYR/PYL family of START proteins. Science 324: 1068–1071

Pertea M, Pertea GM, Antonescu CM, Chang TC, Mendell JT, Salzberg SL (2015) StringTie enables improved reconstruction of a transcriptome from RNA-seq reads. Nat Biotechnol 33: 290–295

Plaxton WC, Tran HT (2011) Metabolic adaptations of phosphate-starved plants. Plant Physiol 156: 1006–1015

Qin T, Zhao H, Cui P, Albesher N, Xiong L (2017) A Nucleus-Localized Long Non-Coding RNA Enhances Drought and Salt Stress Tolerance. Plant Physiol 175: 1321–1336

Raghothama KG (1999) Phosphate Acquisition. Annual Review of Plant Physiology and Plant Molecular Biology 50: 665–693

Robinson D (1994) The responses of plants to non-uniform supplies of nutrients. New Phytol.: 127:635

Rubio V, Linhares F, Solano R, Martin AC, Iglesias J, Leyva A, Paz-Ares J (2001) A conserved MYB transcription factor involved in phosphate starvation signaling both in vascular plants and in unicellular algae. Genes & Development 15: 2122–2133

Santiago J, Rodrigues A, Saez A, Rubio S, Antoni R, Dupeux F, Park SY, Marquez JA, Cutler SR, Rodriguez PL (2009) Modulation of drought resistance by the abscisic acid receptor PYL5 through inhibition of clade A PP2Cs. Plant J 60: 575–588

Schaffer R, Ramsay N, Samach A, Corden S, Putterill J, Carre IA, Coupland G (1998) The late elongated hypocotyl mutation of Arabidopsis disrupts circadian rhythms and the photoperiodic control of flowering. Cell 93: 1219–1229

Secco D, Jabnoune M, Walker H, Shou H, Wu P, Poirier Y, Whelan J (2013) Spatio-temporal transcript profiling of rice roots and shoots in response to phosphate starvation and recovery. Plant Cell 25: 4285–4304

Shiu SH, Bleecker AB (2001) Plant receptor-like kinase gene family: diversity, function, and signaling. Sci STKE 2001: re22

Smith S, Zhu S, Joos L, Roberts I, Nikonorova N, Vu LD, Stes E, Cho H, Larrieu A, Xuan W, Goodall B, van de Cotte B, Waite JM, Rigal A, Ramans Harborough S, Persiau G, Vanneste S, Kirschner GK, Vandermarliere E, Martens L, Stahl Y, Audenaert D, Friml J, Felix G, Simon R, Bennett MJ, Bishopp A, De Jaeger G, Ljung K, Kepinski S, Robert S, Nemhauser J, Hwang I, Gevaert K, Beeckman T, De Smet I (2020) The CEP5 Peptide Promotes Abiotic Stress Tolerance, As Revealed by Quantitative Proteomics, and Attenuates the AUX/IAA Equilibrium in Arabidopsis. Mol Cell Proteomics 19: 1248–1262

Statello L, Guo CJ, Chen LL, Huarte M (2021) Author Correction: Gene regulation by long non-coding RNAs and its biological functions. Nat Rev Mol Cell Biol 22: 159

Stigter KA, Plaxton WC (2015) Molecular Mechanisms of Phosphorus Metabolism and Transport during Leaf Senescence. Plants (Basel) 4: 773–798

Sun N, Wang J, Gao Z, Dong J, He H, Terzaghi W, Wei N, Deng XW, Chen H (2016) Arabidopsis SAURs are critical for differential light regulation of the development of various organs. Proc Natl Acad Sci U S A 113: 6071–6076

Tabata R, Sumida K, Yoshii T, Ohyama K, Shinohara H, Matsubayashi Y (2014) Perception of root-derived peptides by shoot LRR-RKs mediates systemic N-demand signaling. Science 346: 343–346

Thomma BP, Cammue BP, Thevissen K (2003) Mode of action of plant defensins suggests therapeutic potential. Curr Drug Targets Infect Disord 3: 1–8

Thorvaldsdottir H, Robinson JT, Mesirov JP (2013) Integrative Genomics Viewer (IGV): high-performance genomics data visualization and exploration. Brief Bioinform 14: 178–192

Tominaga-Wada R, Wada T (2014) Regulation of root hair cell differentiation by R3 MYB transcription factors in tomato and Arabidopsis. Front Plant Sci 5: 91

Trapnell C, Roberts A, Goff L, Pertea G, Kim D, Kelley DR, Pimentel H, Salzberg SL, Rinn JL, Pachter L (2014) Differential gene and transcript expression analysis of RNA-seq experiments with TopHat and Cufflinks (vol 7, pg 562, 2012). Nature Protocols 9: 2513–2513

Trapnell C, Williams BA, Pertea G, Mortazavi A, Kwan G, van Baren MJ, Salzberg SL, Wold BJ, Pachter L (2010) Transcript assembly and quantification by RNA-Seq reveals unannotated transcripts and isoform switching during cell differentiation. Nat Biotechnol 28: 511–515

Ullah K, Zubia E, Narayan M, Yang J, Xu G (2019) Diverse roles of the E2/E3 hybrid enzyme UBE2O in the regulation of protein ubiquitination, cellular functions, and disease onset. FEBS J 286: 2018–2034

Vlad F, Rubio S, Rodrigues A, Sirichandra C, Belin C, Robert N, Leung J, Rodriguez PL, Lauriere C, Merlot S (2009) Protein phosphatases 2C regulate the activation of the Snf1-related kinase OST1 by abscisic acid in Arabidopsis. Plant Cell 21: 3170–3184

Wada T, Hayashi N, Tominaga-Wada R (2015) Root hair formation at the root-hypocotyl junction in CPC-LIKE MYB double and triple mutants of Arabidopsis. Plant Signal Behav 10: e1089372

Wang Q, Wang J, Yang Y, Du W, Zhang D, Yu D, Cheng H (2016) A genome-wide expression profile analysis reveals active genes and pathways coping with phosphate starvation in soybean. BMC Genomics 17: 192

Wang Y, Lyu W, Berkowitz O, Radomiljac JD, Law SR, Murcha MW, Carrie C, Teixeira PF, Kmiec B, Duncan O, Van Aken O, Narsai R, Glaser E, Huang S, Roessner U, Millar AH, Whelan J (2016) Inactivation of Mitochondrial Complex I Induces the Expression of a Twin Cysteine Protein that Targets and Affects Cytosolic, Chloroplastidic and Mitochondrial Function. Mol Plant 9: 696–710

Wang ZY, Tobin EM (1998) Constitutive expression of the CIRCADIAN CLOCK ASSOCIATED 1 (CCA1) gene disrupts circadian rhythms and suppresses its own expression. Cell 93: 1207–1217

Wilusz JE, Sunwoo H, Spector DL (2009) Long noncoding RNAs: functional surprises from the RNA world. Genes Dev 23: 1494–1504

Woo J, MacPherson CR, Liu J, Wang H, Kiba T, Hannah MA, Wang XJ, Bajic VB, Chua NH (2012) The response and recovery of the Arabidopsis thaliana transcriptome to phosphate starvation. BMC Plant Biology 12: 62

Yong-Villalobos L, Gonzalez-Morales SI, Wrobel K, Gutierrez-Alanis D, Cervantes-Perez SA, Hayano-Kanashiro C, Oropeza-Aburto A, Cruz-Ramirez A, Martinez O, Herrera-Estrella L (2015) Methylome analysis reveals an important role for epigenetic changes in the regulation of the Arabidopsis response to phosphate starvation. Proc Natl Acad Sci U S A 112: E7293–7302

Yu J, Yang L, Liu X, Tang R, Wang Y, Ge H, Wu M, Zhang J, Zhao F, Luan S, Lan W (2016) Overexpression of Poplar Pyrabactin Resistance-Like Abscisic Acid Receptors Promotes Abscisic Acid Sensitivity and Drought Resistance in Transgenic Arabidopsis. PLoS One 11: e0168040

Yuan J, Zhang Y, Dong J, Sun Y, Lim BL, Liu D, Lu ZJ (2016) Systematic characterization of novel lncRNAs responding to phosphate starvation in Arabidopsis thaliana. BMC Genomics 17: 655

Zhu JY, Li Y, Cao DM, Yang H, Oh E, Bi Y, Zhu S, Wang ZY (2017) The F-box Protein KIB1 Mediates Brassinosteroid-Induced Inactivation and Degradation of GSK3-like Kinases in Arabidopsis. Mol Cell 66: 648–657 e644

Bustos R, Castrillo G, Linhares F, Puga MI, Rubio V, Perez-Perez J, Solano R, Leyva A, Paz-Ares J (2010) A central regulatory system largely controls transcriptional activation and repression responses to phosphate starvation in Arabidopsis. PLoS Genet 6: e1001102

Pant BD, Lee S, Lee HK, Krom N, Pant P, Jang Y, Mysore KS (2022) Overexpression of Arabidopsis nucleolar GTP-binding 1 (NOG1) proteins confers drought tolerance in rice. Plant Physiol 189: 988–1004

